# Proteomics of resilience to Alzheimer’s disease identifies brain regional soluble Aβ levels, actin filament processes, and response to injury

**DOI:** 10.1101/2022.10.09.511430

**Authors:** Zhi Huang, Gennifer E. Merrihew, Eric B. Larson, Jea Park, Deanna Plubell, Edward J. Fox, Kathleen S. Montine, C. Dirk Keene, James Y. Zou, Michael J. MacCoss, Thomas J. Montine

## Abstract

Resilience to Alzheimer’s disease (RAD) is an uncommon combination of high disease burden without dementia that may provide critical insights into limiting the clinical impact of this incurable disease. In this study, we used mass spectrometry-based proteomics to quantify regional protein differences that characterize RAD. Starting with over 700 brain donations, we identified 43 extensively annotated research participants who met stringent inclusion exclusion criteria and analyzed matched isocortical regions, hippocampus, and caudate nucleus. Differential expression analysis of 7,115 soluble proteins identified lower isocortical and hippocampal soluble Aβ peptide levels as a significant feature of RAD. Protein co-expression analysis revealed a group of 181 densely-interacting proteins significantly associated with RAD that were enriched for actin filament-based process, cellular detoxification, and wound healing in isocortex and hippocampus. We further support our findings using data from 689 human isocortical samples from four independent external cohorts that were the closest approximations of our clinico-pathologic groups. The molecular basis of RAD, a widely replicated state in older adults for which there is no experimental model, likely holds important insights into therapeutic interventions for Alzheimer’s disease.

## Introduction

Dementia in older individuals is a major medical challenge that looms as a public health disaster unless effective interventions are discovered and deployed [1]. Dementia in older individuals is a syndrome that derives from five different, prevalent diseases. While each of these diseases on its own can cause dementia, in the majority of affected individuals these diseases variably combine in a now widely validated idiosyncratic conspiracy of Alzheimer’s disease (AD), vascular brain injury (VBI), Lewy body disease (LBD), hippocampal sclerosis (HS), and limbic-associated TDP-43 encephalopathy (LATE) [2]. The resulting individually varying comorbidity confounds clinical research because of limited tools to detect each of these five diseases during life; hence, the major focus on developing biomarkers and the continued reliance on brain autopsy to evaluate comprehensively the burden of comorbidities in an individual.

Each of the five commonly comorbid diseases that can contribute to dementia in older individuals has a latent phase. The majority of people harbor a low burden of latent disease that is insufficient to cause dementia, referred to as preclinical [2]. In contrast, a minority harbor a high burden of latent disease(s) sufficient to cause dementia in others; this intriguing group is called resilient, meaning resilient to the clinical expression of dementia despite a sufficiently high burden of disease(s) (https://reserveandresilience.com/framework). Previous proteomic studies have focused on asymptomatic AD (AsymAD), which is a mixture of both preclinical and resilient cases [3–6]. Here, we present the first proteomic study to focus on resilience to AD (RAD).

Latent disease confounds accurate assignment to the control group because without comprehensive neuropathologic assessment the control group will harbor unknown levels of comorbid disease(s) [2]. Comorbidity confounds accurate assignment as RAD; indeed, we have shown in multiple, large population- and community-based cohorts that the major driver of apparent RAD is not related to AD but rather undetected comorbidities that are infrequent in the cognitively resilient group but that are significantly more prevalent in dementia group [7,8]. Here, we have used comprehensive neuropathologic evaluation combined with clinical assessment proximate to death to resolve these confounders and allow accurate clinico-pathologic assignment of both controls free of clinically significant brain diseases and individuals with actual RAD [7,8].

Most proteomic studies have evaluated only one or two isocortical regions that undergo neurodegeneration in AD without including a brain region that does not degenerate to control for coincident events that accompany dementia, like reduced activity and weight loss, that impact the brain but that are thought to be consequences rather than causes of neurodegeneration. Here, we have used data independent acquisition (DIA) MS/MS proteomics of soluble protein extracts from multiple brain regions donated by comprehensively evaluated research participants who were normal controls (NC) free of clinically significant brain diseases, had actual RAD, or had AD dementia (ADD) without significant comorbidities [9]. Our differential expression analysis performed for each brain region identified 33 RAD-associated differentially expressed proteins (DEPs). Protein co-expression analysis revealed a group of 181 densely-interacting proteins that were significantly associated with RAD and enriched for actin filament-based process, cellular detoxification, and wound healing in isocortex and hippocampus. We further supported our findings using data from 689 human isocortical samples from four independent external cohorts that were the closest approximations of our clinico-pathologic groups. The molecular basis of RAD, a widely replicated state in older adults for which there is no experimental model, likely holds important insights into therapeutic interventions for AD.

## Results

Our workflow included four different brain regions (caudate nucleus or CAUD, N=38; hippocampus or HIPP, N=41; inferior parietal lobule or IPL, N=38; and superior and middle temporal gyrus or SMTG, N=38) that were derived from 43 donors, out of 737 brain donations, who met rigorous eligibility criteria for three clinico-pathologic groups: NC (N=11), RAD (N=12), and ADD (N=20) (**Figure 1a, Extended Table 1**, and **Extended Figure 1**). It is important to note that clinically significant co-morbidities were excluded from all groups, NC did not meet consensus criteria for AD, and RAD and ADD were matched for level of AD neuropathologic change (ADNC, P=0.19). Sample preparation and DIA-based proteomics were performed exactly as described [9], and protein level results analyzed by differential expression analysis and co-expression network analysis. Throughout, we compared our results with four independent proteomic datasets whose samples were the closest approximation of our focused study of RAD, including ROS/MAP [10], Banner [11], UPP [12], and BLSA [13] datasets (detailed cohort information is in the **Online Methods** section). A total of 7,115 proteins were quantified among the 155 samples (**Figure 1b**). Corrected Student’s t-test identified 85 significantly differentially expressed proteins (DEPs) among the three clinico-pathologic groups in at least one of the four brain regions, including 33 unique RAD-associated DEPs (RAD DEPs).

**Figure 1.**
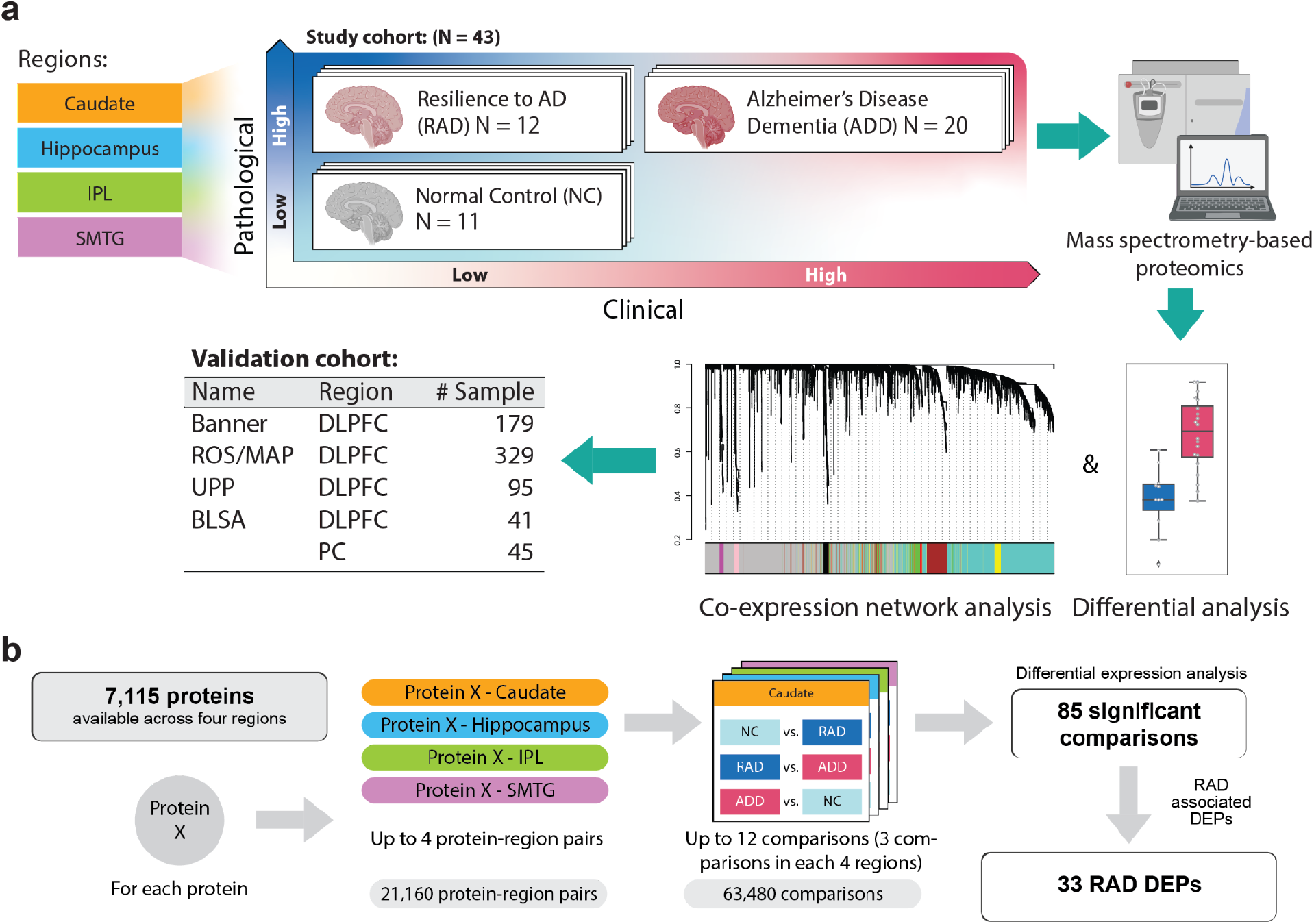
Workflow of this study. **(a)** Samples (N=155) from up to four matched brain regions were donated by 43 research participants who were assigned to three clinico-pathologic groups: NC (normal control), RAD (cognitive resilience to Alzheimer’s disease), or AD dementia (ADD). Samples were quantified by data independent tandem mass spectrometry and data analyzed by differential expression and co-expression network analyses. Results were compared to four independent data sets that most closely approximated our study design. **(b)** Illustration of differential expression analysis and summary of the final number of RAD-associated differentially expressed proteins (RAD DEPs).

### Differential expression analysis

7,115 total proteins were quantified across the four brain regions (**Figure 2a**), of which 5,772 were detected in two or more regions and most proteins (N=3,964) were detected in all four regions (**Figure 2b**). Total proteins first were analyzed by corrected (Benjamini-Hochberg method, FDR cutoff = 0.05) two-sided t-tests, yielding 85 significant protein comparisons between at least one pair of the three groups in at least one brain region (**Figure 2c**). After excluding those DEPs that were not detected in all four regions, we identified 33 unique RAD DEPs (see **Online Methods**) (**Figure 2d, 2e**).

**Figure 2.**
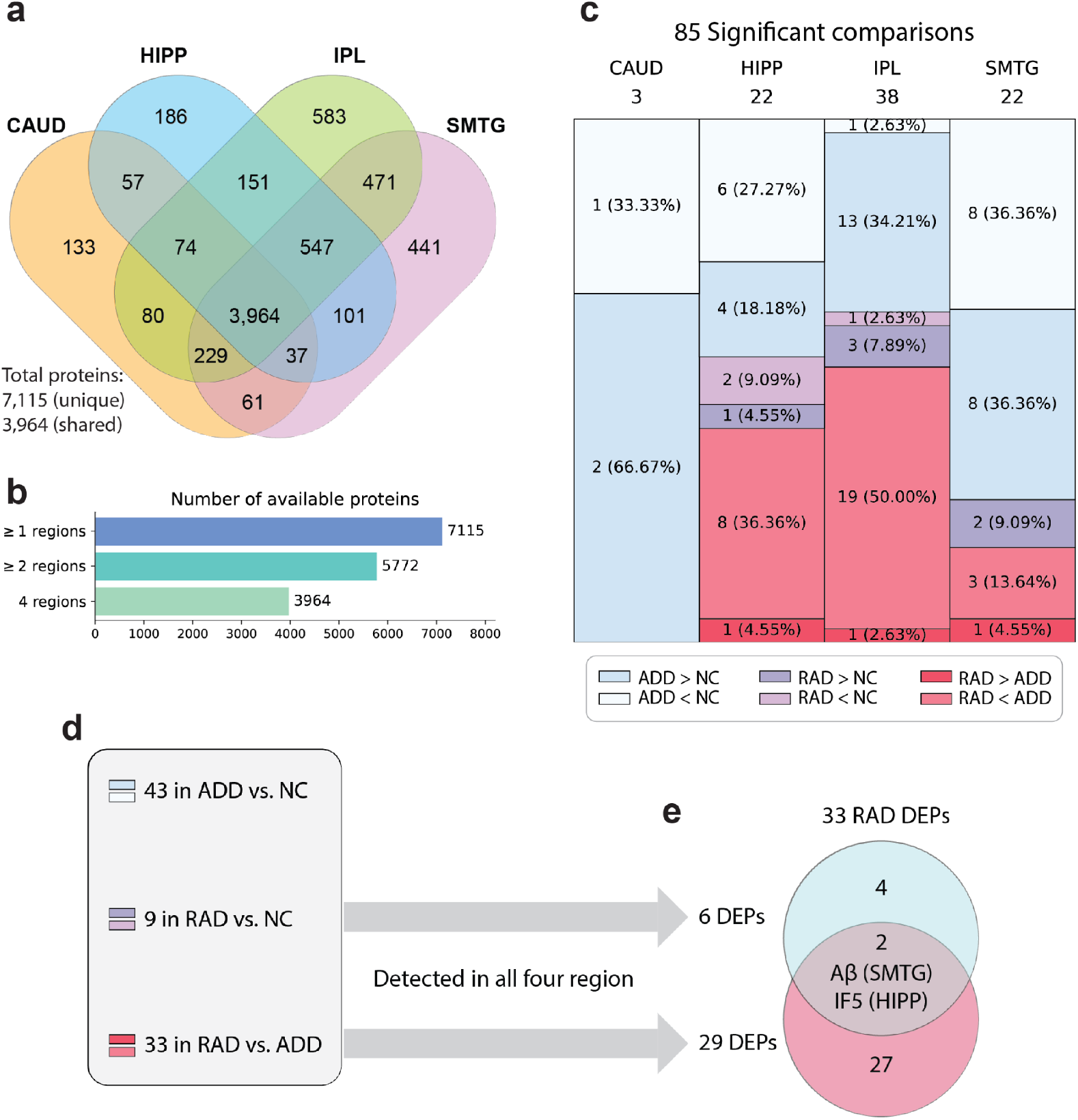
Differentially expressed proteins (DEPs). **(a)** Venn diagram shows overlap in proteins quantified across four brain regions. **(b)** Summary of detected proteins in multiple brain regions. **(c)** 85 proteins were differentially expressed (FDR cut-off = 0.05) among the three clinico-pathologic groups across the four brain regions. **(d)** Of the 85 DEPs, 43 were significantly different between ADD versus NC, and 42 were RAD-associated, meaning significantly different between RAD and either NC or ADD in one or more regions. **(e)** 9 proteins were differentially expressed in RAD versus NC, and 33 proteins were differentially expressed in RAD versus ADD with Aβ and IF5 overlapping. We excluded RAD DEPs that were not detected in all regions, yielding 33 unique RAD DEPs measured in all four regions.

### Regional analysis and validation of RAD DEPs

RAD DEPs were unevenly distributed across brain regions with the most in IPL, then HIPP, and then SMTG (**Figure 3a**). Notably, RAD DEPs were largely non-overlapping across regions with only two RAD DEPs significantly different in more than one region: soluble Aβ in HIPP and SMTG, and CAPG in IPL and SMTG. All RAD DEPs were compared against four independent datasets for validation (**Figure 3b**): Banner Sun Health Research Institute (Banner) [11], Religious Orders Study and Rush Memory and Aging Project (ROS/MAP) [10], UPenn Proteomics study (UPP) [12], and Baltimore Longitudinal Study of Aging (BLSA) [13]. We followed the less stringent case assignment criteria used by others because of limited availability of pathologic data for the external studies and to align with previous publications; the Control (Ctrl), Asymptomatic AD (AsymAD), and ADD groups defined by others were the closest approximation of our more stringently defined groups (**Extended Table 2**), and therefore should provide some level of validation [3,4]. Banner, ROS/MAP, and UPP collected proteomics data in dorsolateral prefrontal cortex (DLPFC), while BLSA collected proteomic data from both DLPFC and precuneus (PC). There was broad agreement among Banner, ROS/MAP, and UPP data; however, BLSA data did not compare well with the other three validation sets, perhaps due to the relatively limited number of samples. Despite these differences in clinico-pathologic criteria and variability in external dataset quality, 70% (23/33) of RAD DEPs in the study set were validated as AsymAD DEPs in at least one external dataset. When limiting the comparison to only isocortical regions in the study set (external dataset exclusively used isocortical regions), 71% of the 24 isocortical RAD DEPs were validated as AsymAD DEPs in one and 58% were validated in two or three external datasets.

**Figure 3.**
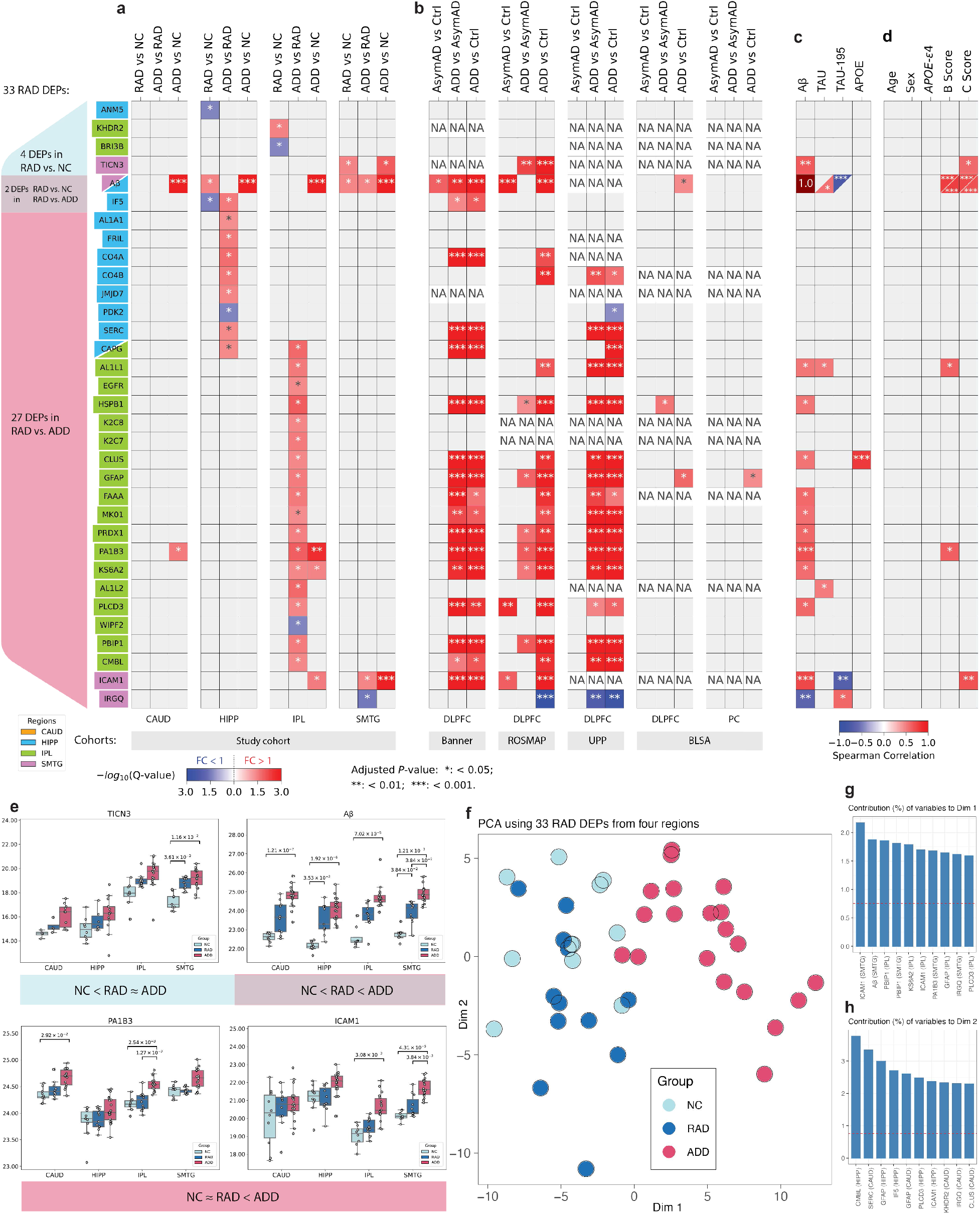
RAD DEPs among four brain regions and four external validation datasets. **(a-b)** Results of corrected multiple comparisons among four brain regions in the study set **(a)**, and the four external validation sets of which only BLSA examined two brain isocortical regions **(b)**. Each column for the study set consists of three comparisons: RAD vs. NC, ADD vs. RAD, and ADD vs. NC. For external datasets, each column consists of three comparisons: AsymAD vs. Ctrl, ADD vs. AsymAD, and ADD vs. Ctrl (non-significant comparisons were colored in gray; insufficient data are white and annotated with “NA”; fold change (FC) < 1 was colored in blue, FC > 1 was colored in red; and colors are the same for -log_10_(Adjusted P-value) ≥ 3). All P-values were adjusted for multiple comparisons (FDR cut-off = 0.05). Annotations: * Adjusted P-value < 0.05; ** Adjusted P-value < 0.01, *** Adjusted P-value < 0.001. **(c-d)** Correlation of expression of each RAD DEP with hallmark AD protein expression in the same brain region **(c)** and with clinical, genetic, or pathologic features of the individual **(d)**. Note: Aβ was a RAD DEP in both HIPP and SMTG, and CAPG was a RAD DEP in both HIPP and IPL. **(e)** Boxplots of selected RAD DEP expression. **(f)** Principal component analysis (PCA) for 33 RAD DEPs in all 4 regions of each brain (original dimension: 33 × 4 = 132) colored by clinico-pathologic groups, visualized in principal dimensions 1 and 2. Variable contributions to the principal dimension 1 **(g)** and principal dimension 2 **(h)** with dashed lines in red showing variable contributions and their expected average.

Regional expression levels were correlated between each RAD DEP and each of four proteins thought to be central to either the etiology or pathogenesis of AD: Aβ, APOE, and TAU as well as the TAU-195 peptide (SGYSSPGSPGTPGSR) that is depleted as TAU is increasingly phosphorylated [14] (**Figure 3c**). Excluding Aβ itself, expression of 12 RAD DEPs was significantly correlated with soluble Aβ concentration; the strongest of these were PA1B3 (P<0.001, for simplicity all “P” stand for “adjusted P-value”) in IPL as well as TICN3 (P<0.01), ICAM1 (P<0.001), and IRGQ (P<0.01) in SMTG; all were positively correlated with Aβ except IRGQ levels in SMTG that were negatively correlated with Aβ levels. Expression of three proteins (Aβ in HIPP, AL1L1 in IPL, and AL1L2 in IPL) were weakly positively correlated with TAU levels in the corresponding region (P<0.05 for each). Aβ (P<0.001) and ICAM1 (P<0.01) levels were negatively correlated with TAU-195 peptide levels in SMTG, indicating that their tissue concentrations increased with increasing tau hyperphosphorylation in this region. IRGQ levels in SMTG were positively correlated (P<0.05) with TAU-195 peptide, suggesting that of all of the RAD DEPs only IRGQ levels in SMTG decreased as both Aβ and hyperphosphorylated tau increased. Only CLUS, also known as apolipoprotein J, in IPL had a significant correlation with APOE protein levels (P<0.001).

Tissue levels of the RAD DEPs were then correlated with individual-level data from each of the 43 donors, and so likely limited to very strong associations. No RAD DEP’s expression correlated significantly with age, sex, or presence of *APOE* ε4 allele (**Figure 3d**). Histopathologic rankings of neurofibrillary degeneration (B score) [15] and neuritic plaque density (C score) [16] were positively correlated with most RAD DEPs that expressed higher in ADD than NC group (**Extended Figure 7**). Aβ levels in HIPP and SMTG were strongly positively correlated with both rankings of neurofibrillary degeneration and neuritic plaque density (P<0.001), aligning well with the Spearman correlations above and confirming our pathological assessments. AL1L1 and PA1B3, both in IPL, were positively correlated with ranking of neurofibrillary degeneration (P<0.05). TICN3 (P<0.05) and ICAM1 (P<0.01) levels in SMTG were positively correlated with ranking of neuritic plaque density. Together, expression levels of five RAD DEPs were significantly correlated with both AD-related protein expression and histopathologic rankings (Aβ in SMTG and HIPP, ICAM1 in SMTG, AL1L1 in IPL, TICN3 in SMTG, and PA1B3 in IPL; **Figure 3d**), while expression of IRGQ in SMTG correlated only with pathologic protein expression.

Finally, we performed principal component analysis (PCA) to inspect higher-level proteomic characteristics by summarizing protein expression levels of the 33 RAD DEPs for all 4 regions into a single value for each individual’s brain (**Figure 3e**). There was broad overlap between NC and RAD groups despite one being free of clinically significant disease and the other having extensive AD neuropathologic change. Furthermore, there was near complete separation of RAD from ADD groups despite both having an equivalent high burden of AD neuropathologic change but only one succumbing to dementia. The top three contributors to PC1 (24.5% of variance) were ICAM1 (SMTG), Aβ (SMTG), and PBIP1 (IPL) (**Figure 3g, Extended Figure 3**), while the top three contributors to PC2 (9.0% of variance) were CMBL (HIPP), SERC (CAUD), GFAP (HIPP) (**Figure 3h, Extended Figure 3**).

### Protein co-expression network analysis

To nominate related proteins that robustly can distinguish RAD, we expanded the analysis from individual proteins to protein modules using the established WGCNA algorithm [17] to perform a consensus weighted protein co-expression network analysis on the 3,964 proteins detected in all four regions (**Figure 4a**). The resulting 9 co-expression modules were then used to estimate eigenproteins, which can be considered as the summary of a module’s overall protein expression [18]. As expected, the two isocortical regions had similar eigenprotein expression compared to the other two regions. The regions that undergo neurodegeneration in AD, HIPP, IPL, and SMTG, but not CAUD, showed significant positive correlations between clinico-pathologic groups and module (M) 5, while M1 in HIPP was negatively correlated with clinico-pathologic groups (**Figure 4a**). Among individual-level data including age, sex, B score, C score, and *APOE* ε4 allele, only age was positively correlated with M1 in both IPL and SMTG and negatively correlated with M4 in both IPL and SMTG (**Extended Figure 5**). The 33 RAD DEPs were distributed across M0, M1, and M5 (**Figure 4b**). Specifically, M0 contained Aβ, AL1L1, AL1L2, ANM5, EGFR, IF5, JMJD7, KHDR2, MK01, TICN3, and KS6A2; M1 contained BRI3B, AL1A1, FRIL, PDK2, SERC, PLCD3, WIPF2, and PBIP1; and M5 contained C04A, C04B, CAPG, HSPB1, K2C7, K2C8, CLUS, GFAP, FAAA, PRDX1, PA1B3, CMBL, ICAM1, and IRGQ. We performed Spearman correlation analysis for the 33 RAD DEPs within each region and summed the number of significantly correlated (Adjusted P-value < 0.05) proteins across all regions (**Extended Figure 6**). Of the 13 least correlated proteins, 11 of them were assigned to M0, including Aβ.

**Figure 4.**
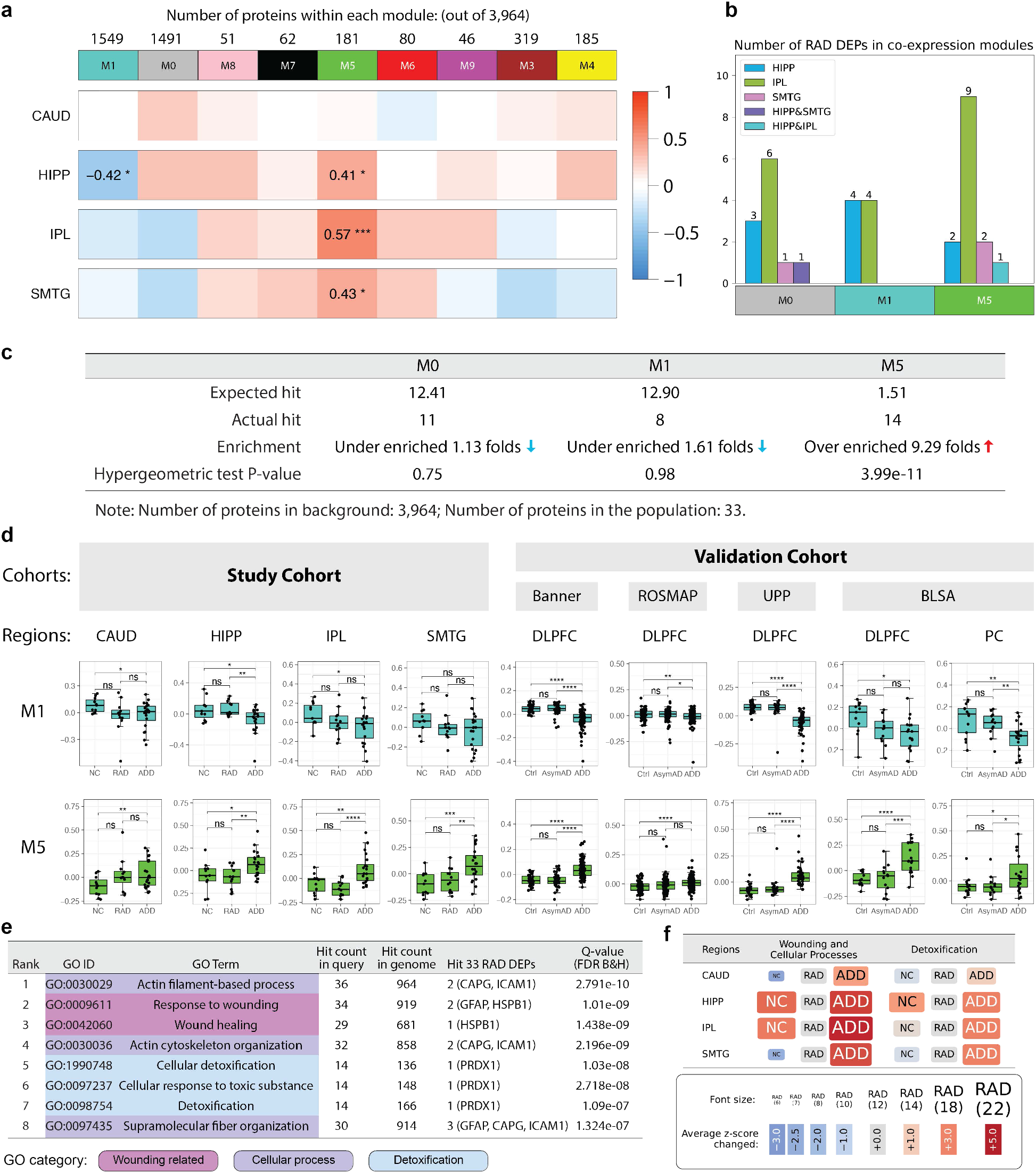
**(a)** Consensus protein co-expression analysis identified 9 modules across four brain regions. Bi-weighted mid-correlation was used to evaluate the relationships between clinico-pathologic groups and eigenprotein expression (correlation text threshold: ±0.4). All P-values were adjusted by the Benjamini-Hochberg procedure (FDR cut-off = 0.05). **(b)** Three co-expression modules contained the 33 RAD DEPs. **(c)** The number of expected and observed RAD DEPs in each module, and enrichment analysis via hypergeometric test. **(d)** Module 1 and 5 eigenprotein expressions in NC, RAD, and ADD for the study set and in Ctrl, AsymAD, and ADD for external validation sets. **(e)** Top 3 enriched GO biological process categories in M5 and their enrichment analysis results. **(f)** Patterns of the change in M5 z-scores. Font sizes of clinico-pathologic groups reflect average z-score changes within the GO categories.

A hypergeometric test evaluated the expression of the RAD DEPs among the consensus AD network modules (**Figure 4c**). M5 was strongly enriched for expression of RAD DEPs (14 of 33 RAD DEPs; P-value = 3.99e-11), while M0 and M1 were not significantly enriched in RAD DEPs (M0 P-value = 0.75, M1 P-value = 0.98). Indeed, M5 was over-enriched in RAD DEPs by almost 10-fold compared to chance. M5 eigenprotein expression was then compared among the study set and external datasets (**Figure 4d**). We found the same dementia-associated pattern (NC≈RAD<ADD) across all regions and all datasets, robustly validating that M5 eigenprotein expression was significantly greater in ADD compared to RAD or NC groups or compared to AsymAD or Ctrl groups. Furthermore, a protein-protein interaction (PPI) network for M5 was constructed based on the STRING database v11.5 [19]. Among the 181 proteins in M5, 177 primary genes were identified in the PPI network with 468 edges in total (average node degree = 5.29), indicating that PPI in M5 has significantly more interactions than expected by chance (P < 1.0e-16). **Extended Figure 8** shows the PPI network of M5 with experimentally determined interactions highlighted. Together, these results underscore that M5 is a robustly validated and densely co-expressed module of 181 proteins that distinguishes ADD from RAD despite their equivalent histopathologic burden of disease (**Extended Figures 11-15**).

### Enrichment analysis of M5

We performed gene ontology (GO) analysis using each M5 protein’s primary gene [20] and identified the three top GO categories based on the branches in the ancestor chart: (i) Wounding Related (GO:0009611, GO:0042060), (ii) Cellular Process (GO:0030029, GO:0030036, GO:0097435), and (iii) Detoxification (GO:1990748 GO:0097237 GO:0098754). These top three GO categories contained the eight strongest GO terms according to enrichment P-values (**Figure 4e**) and overlapped with sixty-nine of M5 proteins’ primary genes (**Extended Figure 9b**). Among these GO terms, Actin Filament-based Process (GO:0030029, Q-value = 2.791e-10) ranked highest with its descendant Actin Cytoskeleton Organization (GO:0030036, Q-value = 2.196e-09) ranked fourth. These two plus Supramolecular Fiber Organization (GO:0097435, Q-value = 1.324e-07) were grouped into the Cellular Process GO category (light purple) and shared two RAD DEPs: ICAM1 and CAPG. The Wounding Related GO category (dark purple), including Response to Wounding (GO:0009611, Q-value = 1.01e-09) and its subcategory Wound Healing (GO:0042060, Q-value = 1.438e-09), shared the RAD DEP HSPB1. The Detoxification GO category (light blue), including Cellular Response to Toxic Substance (GO:0097237, Q-value = 2.718e-08), Detoxification (GO:0098754, Q-value = 1.09e-07) and its descendant Cellular Detoxification (GO:1990748, Q-value = 1.03e-08) contained the RAD DEP PRDX1 (**Figure 4e**). Finally, a few regional and group patterns emerged when we calculated z-scores to assess whether the items in the different GO terms were up- or down-regulated (**Figure 4e, Extended Figure 10**). Wounding Related and Cellular Process are grouped because they always changed together with a consistent pattern of RAD several fold less than ADD in HIPP and isocortical regions and less pronounced group differences in CAUD. Detoxification repeated this general expression pattern with strong increase in ADD compared to RAD in HIPP and isocortical regions with less pronounced differences among groups in CAUD. In all GO categories for all regions, RAD more closely approximated NC than ADD (**Figure 4f**).

## Discussion

Our study focused on actual resilience to the clinical expression of dementia from AD, and because of this it had important design differences from previous proteomic studies of AD. [3] [21] [22] [4]. The most important difference is that we comprehensively evaluated brains to minimize confounding from latency in NC and from comorbidities in all groups. Exclusion of latent and comorbid diseases had the inevitable consequence of reducing the number of cases eligible for study; indeed only 43 of 737 eligible brain donations met our stringent criteria. In part to offset the impact of a relatively small number of high quality cases and controls, we expanded our study to include multiple brain regions involved or uninvolved by neurodegeneration. Our approach only included intermediate or high ADNC in the matched RAD and ADD groups (P=0.19), all with sufficient burden of AD to cause dementia (see **Extended Table 3**) [1,40]. In contrast to the approach used by others, our focus on RAD excluded the large number of preclinical AD cases from our cohort; others instead have analyzed AsymAD, which is a mix of preclinical AD and RAD [3,4]. As far as we are aware, our focus on RAD is a unique study design.

The strongest and most consistent RAD signal from our multiregional analysis was soluble Aβ expression, which tended to be in between NC and ADD in all regions, and uniquely Aβ expression in SMTG was significantly less in NC than RAD which in turn was significantly less than ADD. Indeed, soluble Aβ expression in SMTG was a major contributing variable to near complete separation of NC/RAD from ADD using PCA. These results suggest that although RAD and ADD groups were matched by the admittedly coarse tools for histopathologic scoring of ADNC, which includes ordinal ranking of insoluble Aβ plaques, lower tissue concentration of soluble Aβ might be a significant molecular feature of RAD. However, we recognize that our histopathologic matching is imperfect, and it is possible that the decreased soluble Aβ in RAD compared to ADD might be related in part to statistically insignificant variation in overall Aβ levels between these two groups. The soluble, lower molecular mass assemblies of Aβ are typically extracted with SDS (as done here) or urea, while extraction of higher molecular weight, insoluble aggregates of Aβ requires stronger chaotropics, like 70% formic acid [23]. Banner and ROSMAP proteomic data for AsymAD, which contains both preclinical and RAD cases, validated our major finding that aligns readily with abundant experimental data showing that low molecular weight soluble aggregates of Aβ, forms not detected by histopathologic and PET methods that visualize insoluble Aβ fibrils, are concentration-dependent direct neuronal stressors and indirect neuronal stressors via glial cell activation [24,25]. Together, these cross-validating results support that lower tissue concentration of soluble Aβ in isocortex may be a molecular feature of RAD, plausibly resulting in reduced neuronal stress and injury. Most therapeutic antibodies that reduce brain Aβ target larger, less soluble forms of Aβ; so far these have largely failed in clinical trials [26]. In contrast, emerging agents like mAb158, also called BAN2401 or Lecanemab, binds to soluble Aβ species and has shown promising outcomes in initial clinical trials (https://www.alzforum.org/therapeutics/lecanemab) [27]. Our quantitative results from people resilient to ADD could provide a rough estimate of the extent to which soluble Aβ may need to be lowered in isocortical regions to suppress the clinical expression of severe cognitive impairment along the continuum of AD [28].

We pursued other potential contributors to RAD through co-expression network analysis and observed that one module, M5 in HIPP, SMTG, and especially IPL, was significantly enriched in ADD resilience-associated proteins, a result validated in DLPFC using Banner and UPP, and both regions using BLSA data. Across our study and all validation sets, M5 had the pattern NC≈RAD<ADD, implying that increased expression of M5 proteins may be a molecular signature in isocortical and HIPP regions of progression to dementia. In contrast, M1 in HIPP was significantly underrepresented in RAD (NC≈RAD>ADD), a pattern validated in most external datasets and perhaps a consequence of the extensive neurodegeneration in this region that accompanies progression to dementia. Interestingly, Aβ was not a component of M5 or M1 but rather M0, meaning that it was not contained within a co-expression network despite its increasing tissue concentration being a distinguishing feature between RAD and ADD.

Functional insight into the 181 component proteins of strongly resilience-associated M5 by GO analysis yielded the three top categories including actin cytoskeleton organization, wound healing, and cellular detoxification. Actin filament dynamics are essential to dendrite and synapse formation and remodeling, and others have identified actin filament-based processes in enrichment analyses of AD [29] and bipolar disorder [30]. Importantly, preserved synaptic density is a feature of resilience to AD, potentially linking our proteomic data with morphological data to highlight synaptic plasticity as a key compensatory feature of resilience [31]. Wound healing and detoxification are complex responses to injury that showed a similar pattern of expression as actin filament dynamics such that these three biological processes, which represent compensatory change and response to injury, were several fold greater in ADD than RAD in the three regions undergoing neurodegeneration. The lower response to injury in HIPP and isocortical regions of RAD aligns well with our results showing lower soluble Aβ concentration in these regions as well as with the extensively described mechanisms of Aβ induced injury to neurons through both direct and indirect mechanisms [32,33]. Together, these results suggest that RAD is a state of equivalent histopathologic features but lower soluble Aβ and less injury than ADD.

Of the 33 RAD DEPs identified, M5 contained 14 and a few of these deserve specific mention. CLUS, or clusterin, is also known as apolipoprotein (apo) J. Variants in the CLUS gene have been repeatedly associated with the risk of AD (http://www.alzgene.org/). Like the closely functionally related APOE protein whose concentration was strongly positively correlated, CLUS plays a major role in lipid transport in brain where it critically supports synaptic remodeling and repair, and modulates innate immune responses involving the RAD DEPs CO4A and CO4B. ICAM1 is a master regulator of inflammation and injury resolution and was a major contributing variable in multiple brain regions to the near complete separation of NC/RAD from ADD by PCA [34]. Immune-mediated neuronal injury is widely supported as a major contributor to AD pathogenesis, and variants in the gene encoding ICAM1 have been associated with risk of AD (http://www.alzgene.org/) [32]. The platelet-activating factor acetylhydrolase isoform 1B complex, of which PA1B3 is a subunit, is broadly expressed across neuronal cell types and is critical in human brain development, including neuron migration [35,36]. PA1B3 was one of only four RAD DEPs whose concentration increased with both Aβ and hyperphosphorylated tau. Also known as PAFAH1B3, this protein was recently shown by elegant proteomic work of others investigating Ctrl, AsymAD, and ADD samples from ROSMAP to reside at the center of a MAPK/metabolism module of proteins associated with both amyloid plaques and neurofibrillary tangles [3].

Intriguing anatomical differences also were observed in our data. The CAUD subserves motor, cognitive, and behavioral functions, and has been proposed as a site for temporary functional compensation in AD [37,38]. Neither our differential expression analysis nor consensus protein expression analysis identified significant changes in CAUD. Rather both types of analysis localized proteomic changes of RAD to HIPP and isocortex, further suggesting that proteomic changes associated with RAD are unlikely to be nonspecific brain changes that accompany systemic impacts of dementia. The two isocortical regions investigated, SMTG and IPL, had only one overlapping RAD DEP, CAPG. Murine *capg* expression was shown recently to be among a small set of disease-associated microglia genes uniquely upregulated by APOE4 in preclinical models of AD [39]. Resilience-associated proteomic changes were largely non-overlapping in IPL or SMTG, and were only partially validated in prefrontal cortex or precuneus from external datasets, perhaps indicating regional variation in isocortical contributions to RAD. Finally, when comparing up or down regulation of Wounding Related and Cellular Process GO terms, CAUD and SMTG displayed the intriguing pattern of down regulation in RAD vs. NC and upregulation in RAD vs. ADD, while HIPP and IPL showed upregulation in both group comparisons. Although the significance of these region-by-group interactions are not clear, they underscore the likely regional variation in molecular mechanisms of RAD.

This study has several limitations. First, because we used stringent criteria for cohort enrollment, a relatively small group of 43 high-quality cases out of 737 eligible was assembled to focus on RAD. Second, bulk tissue proteomics lacks cell-type specificity and likely obscures important cell-specific changes. In light of our results, we hope future studies will focus on single-cell analysis to further disclose cell type-specific changes in RAD. Lastly, restricted by the availability of stringently defined cases, we conducted independent validation using external datasets that most closely approximated ours. We hope our results motivate others to try to validate our findings by creating an animal model or developing similarly stringently defined clinico-pathological cohorts.

In conclusion, we have undertaken a novel proteomic analysis of carefully annotated human brain regions to determine molecular features of RAD. When compared to ADD, our validated results show that lower tissue concentration of soluble Aβ in isocortical regions as well as lower expression of actin filament-based processes and cellular detoxification/repair in isocortex and hippocampus are characteristic of RAD. Combined with the results of others, our study suggests that people with RAD have lower disease-specific injury, perhaps from less soluble Aβ, and thereby an appropriately limited response to injury. These results provide critical insights into the molecular features of RAD and suggest potential therapeutic strategies to limit the clinical progression to dementia of this increasingly prevalent and incurable disease.

## Acknowledgments

This work was supported by NIH grants AG053959, AG006781, AG066509, and AG066567, and the Nancy and Buster Alvord Endowment.

## Online Methods

### Clinico-pathologic groups

This study was approved by the Institutional Review Boards of the University of Washington and Stanford University. Cognitive diagnosis of dementia or not dementia was made using DSM-IVR criteria; an initial provisional diagnosis of dementia was followed one year later with a confirmed diagnosis of dementia. Out of a total of 340 research brain donations that had been dissected and flash frozen within 8 hours (mean ± SD = 4.4 ± 1.3 hours) of death, 43 cases met eligibility criteria for rigorously defined clinico-pathologic groups (**Supplementary Table 1**): (i) Normal controls (NC, N=11) had neuropsychological test results in the upper quartile for the cohort at their last visit within 2 years of death, did not have AD neuropathologic change (ADNC) according to NIA-AA guidelines, and had clinically insignificant (none/low) pathologic changes of VBI, LBD, HS, or LATE [1,40,41]; (ii) Cognitive resilient to AD (RAD, N=12) had neuropsychological test results in the upper quartile for the cohort at their last visit within 2 years of death, had intermediate or high level ADNC according to NIA-AA guidelines, and had none/low pathologic changes of VBI, LBD, HS, or LATE [1,40,41]; and (iii) AD dementia (ADD, N=20) were diagnosed with dementia during life, had intermediate or high level ADNC according to NIA-AA guidelines, and none/low had pathologic changes of VBI, LBD, HS, or LATE [1,40,41]. Importantly, we excluded from all clinico-pathologic groups cases with low-level ADNC according to NIA-AA guidelines [1,40]. We have shown previously insignificant interval change in diagnosis of not dementia over 2 years between last research evaluation and death for individuals who had neuropsychological test results in the upper quartile for the cohort [42]; all NC and RAD participants had < 2 years between last evaluation and death with an average interval of 352 days. Criteria for including only none/low levels of the four prevalent comorbidities that do not significantly contribute to the risk of dementia were: (i) for VBI: no territorial or lacunar infarcts, no hemorrhages, <2 microinfarcts/microhemorrhages [43], (ii) for LB: none or amygdala only [44], (iii) for LATE-NC: none or amygdala only [45], and (iv) no hippocampal sclerosis in the unilateral hippocampus available for histopathologic analysis.

### Sample preparation and proteomic analysis

Two 25 μm thin frozen sections from the four brain regions were maintained at -80 °C until protein extraction with SDS and DIA proteomic analysis as previously described [9]: caudate nucleus (CAUD, N=38), hippocampus (HIPP, N=41), inferior parietal lobule (IPL, N=38), and superior and middle temporal gyrus (SMTG, N=38) for a total of 155 samples (30 cases were matched across all four regions; **Extended Figure 1**).

The Skyline documents, raw files for quality control and DIA data are available at Panorama Public https://panoramaweb.org/ADBrainCleanDiagDIA.url. ProteomeXchange ID: PXD034525. Access URL: http://proteomecentral.proteomexchange.org/cgi/GetDataset?ID=PXD034525. Metadata is available in **Supplementary Table 1**.

Peptide data was acquired by DIA mass spectrometry (DIA-MS) based on the method we previously described [46]. Protein expression was calculated by aggregating peptide abundances. After *log*_2_(*x*) transformation, median normalization was performed to adjust for minor sources of variability that are difficult to either control or predict, followed with batch correction to remove the effect of different experimental batches. In external datasets, for proteins with two or more isoform identifiers, we kept the expressions with maximum read counts.

### Statistical analysis

All protein expressions were *log*_2_(*x*) transformed before statistical analyses. For differentially expressed proteins analysis, two-sided Student’s t-tests were performed with P-values adjusted for multiple comparisons. For correlation analysis, Spearman’s rank correlation method was performed to evaluate the protein expressions. In consensus weighted protein co-expression analysis, clinico-pathological groups and other factors given module eigenproteins was analyzed by bi-weighted mid-correlation [47]. For analyzing the concentration of RAD DEPs among co-expressed modules and calculating Q-values in gene ontology enrichment analysis [20], hypergeometric test [48] with probability mass function

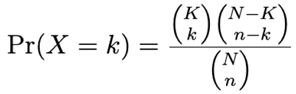

was adopted, where the binomial coefficient is defined as

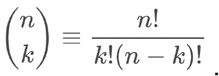

In above equation, *N* is the population size (total number of genes in the background), *K* is the number of hits in the targeted gene list, *n* is the number of draws in the protein list of interests, and *k* is the number of observed successes (The total number of mutual proteins in both protein list of interests and the targeted protein list).

For low-dimensional visualization, principal component analysis (PCA) was adopted.

### Derivation of 33 RAD DEPs

Of the 85 significant protein comparisons (**Figure 2c**), 76% (n=65), were unique indicating that most group differences in protein expression were region-specific. The regional distribution of the 85 DEPs was coded for significant paired group differences: 43 had significantly different expression between ADD and NC, including only 3 in CAUD (Aβ, MT3, PA1B3) with the Aβ result in ADD CAUD confirming our group assignments. Importantly, the 42 preliminary RAD DEPs with significantly different expression between RAD and NC or between RAD and ADD were restricted only to HIPP and isocortical (IPL and SMTG) regions. Nine of the 42 preliminary RAD DEPs had significantly different expression when comparing RAD vs. NC (3 RAD<NC and 6 RAD>NC), and 33 had significantly different expression when comparing RAD vs. ADD (3 RAD>ADD and 30 RAD<ADD). With Aβ and CAPG expression significantly different in two regions, and Aβ in SMTG and IF5 in HIPP overlapping between RAD vs. NC and RAD vs. ADD, there were 38 unique preliminary RAD DEPs. Five of the 38 unique preliminary RAD DEPs were not detected in all four regions (**Figure 3a**); these five were excluded from further analysis, yielding 33 RAD DEPs for detailed analysis.

### Handling missing values

When performing univariate analysis, proteins for which all expressions are missing were removed. If a protein has less than 3 available regional sample expressions, the Student’s t-test and Spearman correlation will not be performed for that protein (use missing value instead). When performing low dimensional visualization and consensus protein co-expression analysis, missing protein expressions were imputed by mean values from other individuals within a certain brain region. When validating protein co-expression modules with external cohorts, proteins were either discarded if all values were unavailable or imputed by mean values from other individuals if available.

### Consensus protein co-expression module analysis

For consensus protein co-expression module analysis, we adopted the WGCNA algorithm and chose soft power = 7 according to **Extended Figure 4b**. And by default, we chose deep split = 2, minimum module size = 30, and merging cut height = 0.25. The consensus co-expression module analysis takes all four brain regions into account, and developed a single set of modules. After consensus merging, module 2 was merged into module 1, and module 0 represents a group of unassigned proteins, resulting 9 modules. Based on categorical groups conditions (NC: 0, RAD: 1, ADD: 2), bi-weighted mid-correlation [47] was performed to evaluate the relationships between clinico-pathologic groups and expression of the 9 eigenproteins, followed with P-value adjustment (FDR cut-off = 0.05).

### Gene ontology enrichment analysis

Gene ontology (GO) enrichment analysis was carried out by ToppGene suite (version 2022-03-28. 20,669 genes in category) [20]. Protein’s primary gene was derived according to UniProtKB (2022_01) database. Ancestor chart was constructed according to AmiGO2 Gene Ontology database [49]. Z-scores were calculated based on number of up/down regulated proteins from hit count in query:

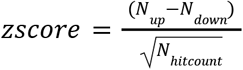

Up and down regulation was based on log2 fold change. If a z-score > 0 on RAD vs NC, then the associated GO term is more likely to be increased when RAD > NC [50]. We set z-score = ±1 as the cut-off threshold to determine the up/down regulation signal of a GO term. Patterns of the change in M5 z-scores were described in fontsize = 12 + 2 × (*zscore*_*avg*_), where *zscore*_*avg*_ is the average change of z-score within a GO category when comparing RAD to other clinico-pathologic groups.

### Validation with External Datasets

For external validation, 179 individuals (Ctrl=42, AsymAD=45, ADD=92) were collected from Banner Sun Health Research Institute (Banner) [11] in dorsolateral prefrontal cortex region (DLPFC, Brodmann area 9), followed with batch effects removal via ComBat [51]. 329 individuals (Ctrl=78, AsymAD=89, ADD=162) were collected from Religious Orders Study and Rush Memory and Aging Project (ROS/MAP) [10] in DLPFC region (TMT quantitation, version: 03/22/2022, SwissProt and TrEMBL human protein db 2015, median polish corrected relative reporter abundance, followed with *log*_2_(*x*) transformation), and 95 individuals (Ctrl=26, AsymAD=20, ADD=49) were collected from the UPenn Proteomics study (UPP) [12] in DLPFC region (label-free quantitation, version: 03/22/2022, median polish ratio over global internal standard (GIS), batch corrected relative reporter abundance, followed with *log*_2_(*x*) transformation). In addition, 41 individuals (Ctrl=11, AsymAD=13, ADD=17) in DLPFC region and 45 individuals (Ctrl=13, AsymAD=13, ADD=19) in precuneus region (PC, Brodmann area 7) were collected from Baltimore Longitudinal Study of Aging (BLSA) [13].

Although these external datasets were the most closely related, there were potentially important differences in clinico-pathologic group assignments between our study and these four external datasets.

1. Controls: By expert consensus guidelines from NIA-AA, our NC group was free of ADNC and clinically significant levels of the four other commonly comorbid diseases, while the approach used by the external datasets permitted low level ADNC in the control group and did not exclude VBI, LBD, HS, and LATE-NC from the control group.
2. RAD vs. AsymAD: Our approach only included intermediate or high ADNC in the RAD group, while the approach used by others permitted low level ADNC in the AsymAD group (preclinical AD) [3,4]. Intermediate or high level ADNC is sufficient to cause dementia [1,40], meaning that our approach focused on RAD while AsymAD is a mix of preclinical AD and RAD cases (see **Extended Table 2**).
3. Apparent vs. actual resilience: Neuropathologic assessment of VBI, LBD, HS, and LATE-NC was not included in the earlier proteomic studies, so these diseases not only are unknowingly present in the control group (*vide supra*), but apparent resilience cannot be distinguished from actual RAD without evaluation of all five diseases [8].
4. Dementia was classified differently among all datasets: for ROSMAP, dementia was clinical cognitive diagnosis summary at last visit (dcfdx_lv) >1 [3,4], which includes mild cognitive impairment and dementia; for Banner, dementia was last MMSE < 24 [3,4]; for BLSA and UPP dementia was “AD” diagnosis code [3,4].

Our more stringent criteria fall within those used by others, meaning that our NC was a subset of external Ctrl, our RAD was a subset of external AsymAD, and our ADD was a subset of external ADD. When we applied our more stringent criteria for ADNC to the external datasets, between about one-quarter and one-half of cases were excluded. If the other four comorbidities had been evaluated, then even more external cases and controls would have been excluded, underscoring the reality of the limited availability of high-quality samples to investigate RAD. We struck the balance of comparing our results for NC, RAD, and ADD to Ctrl, AsymAD, and ADD of the most closely related external datasets as the best available external validation.

## Supplementary Information

Supplementary table is available along with the manuscript.

## Reporting Summary

### Ethics oversight

All study cohort participants were collected and provided informed consent under protocols approved by the Institutional Review Board (IRB) at University of Washington and Stanford University.

## Extended Tables and Figures

**Extended Table 1.** Characteristics of study participants and samples. Complete metadata is listed in **Supplementary Table 1**.

|  |  | Clinico-pathologic groups |  |  | Total |
| --- | --- | --- | --- | --- | --- |
|  |  | NC | RAD | ADD |  |
| Cohort summary | Unique number of individuals | 11 | 12 | 20 | 43 |
|  | Unique number of samples | 37 | 44 | 74 | 155 |
|  | Age (yr; median, min-max) | 88 (73 - 95) | 90 (78 - 98) | 85 (59 - 100) | 88 (59 - 100) |
|  | Sex (% Male) | 54.55% | 50.00% | 60.00% | 55.81% |
| Brain regions | CAUD (number of samples) | 10 | 10 | 18 | 38 |
|  | HIPP (number of samples) | 10 | 11 | 20 | 41 |
|  | IPL (number of samples) | 8 | 12 | 18 | 38 |
|  | SMTG (number of samples) | 9 | 11 | 18 | 38 |
|  |  | Number of proteins |  | Number of peptides |  |
| Regional proteomics data | CAUD | 4,635 |  | 26,123 |  |
|  | HIPP | 5,117 |  | 31,863 |  |
|  | IPL | 6,099 |  | 42,492 |  |
|  | SMTG | 5,851 |  | 40,342 |  |
|  | Total | 7,115 (unique), 3,964 (shared) |  | 56,073 (unique), 19,218 (shared) |  |

Abbreviations: NC for normal control, RAD for resilience to AD, ADD for Alzheimer's disease and dementia, CAUD for Caudate, HIPP for Hippocampus, IPL for Inferior parietal lobule, and SMTG for superior and middle temporal gyri. NC for normal control, RAD for cognitive resilience to Alzheimer's disease, and ADD for Alzheimer's disease dementia.

**Extended Table 2.** The criteria for clinicopathologic groups for the current study and datasets used for external validation are summarized. Note that unlike the external validation datasets, the current study explicitly excluded*: cases with any LB or LATE-NC other than amygdala, >2 microinfarcts or microhemorrhages, any territorial or lacunar infarcts, any hemorrhages, any HS, and any other neuropathologic features of the disease.

| Clinico-pathologic groups |  | Clinical |  |  |  |
| --- | --- | --- | --- | --- | --- |
|  |  | Dementia: No |  | Dementia: Yes |  |
|  |  | Current Study | External Validations | Current Study | External Validations |
| Pathological | Not ADNC | Normal Control (NC)* | Control (Ctrl) | Excluded<br>(Causes of dementia other than AD) |  |
|  | Low ADNC | Excluded<br>(Preclinical AD) |  |  |  |
|  | Intermediate or High ADNC | Resilient to AD (RAD)* | Asymptomatic (Asym) AD | ADD* | ADD |

**Extended Table 3.**
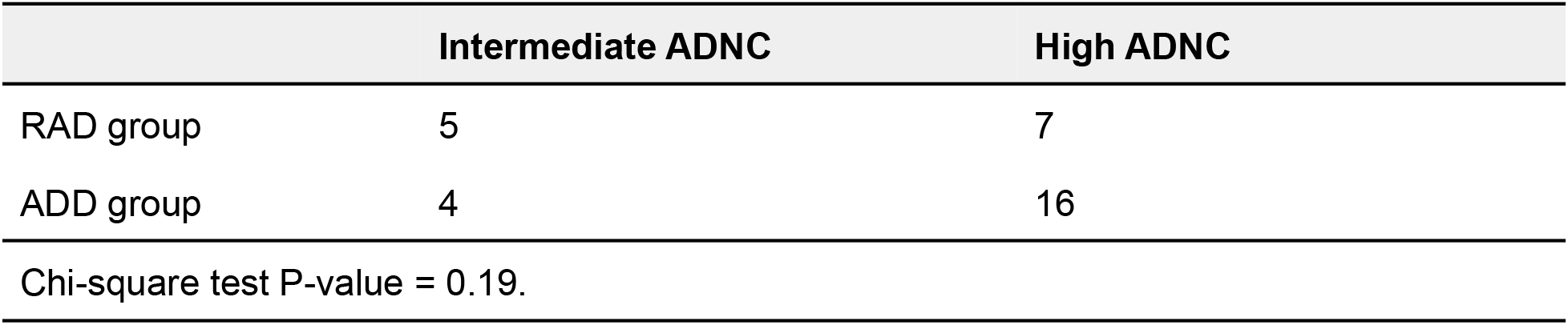
The Chi-square test compares the intermediate and high Alzheimer’s disease neuropathologic change (ADNC) between RAD and ADD groups.

**Extended Figure 1.**
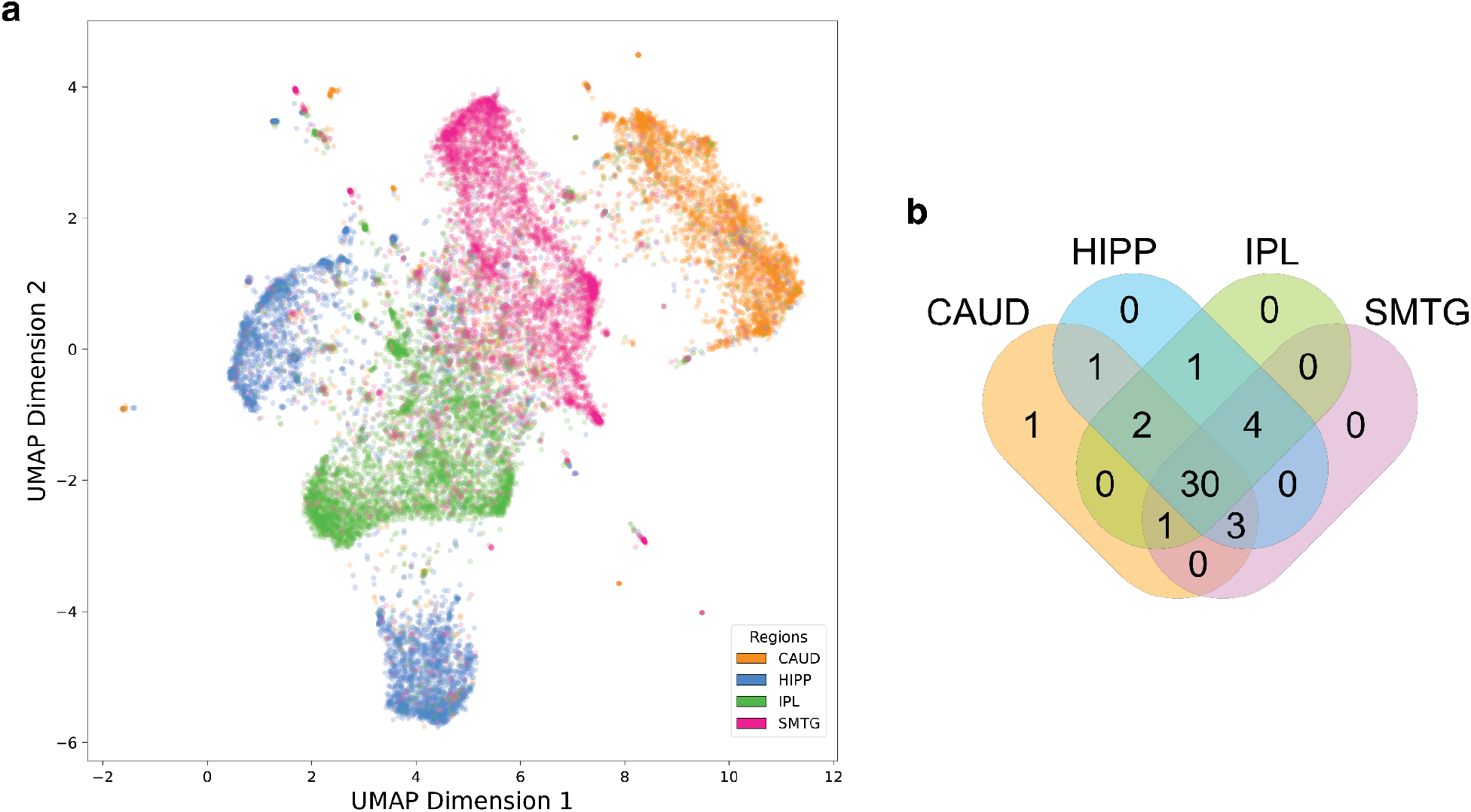
Additional proteomic features. (A) UMAP plot of detected protein expressions across four brain regions from the 43 individuals. (B) Venn diagram of available individuals among four brain regions. 30 out of 43 individuals had proteomic data across four brain regions.

**Extended Figure 2.**
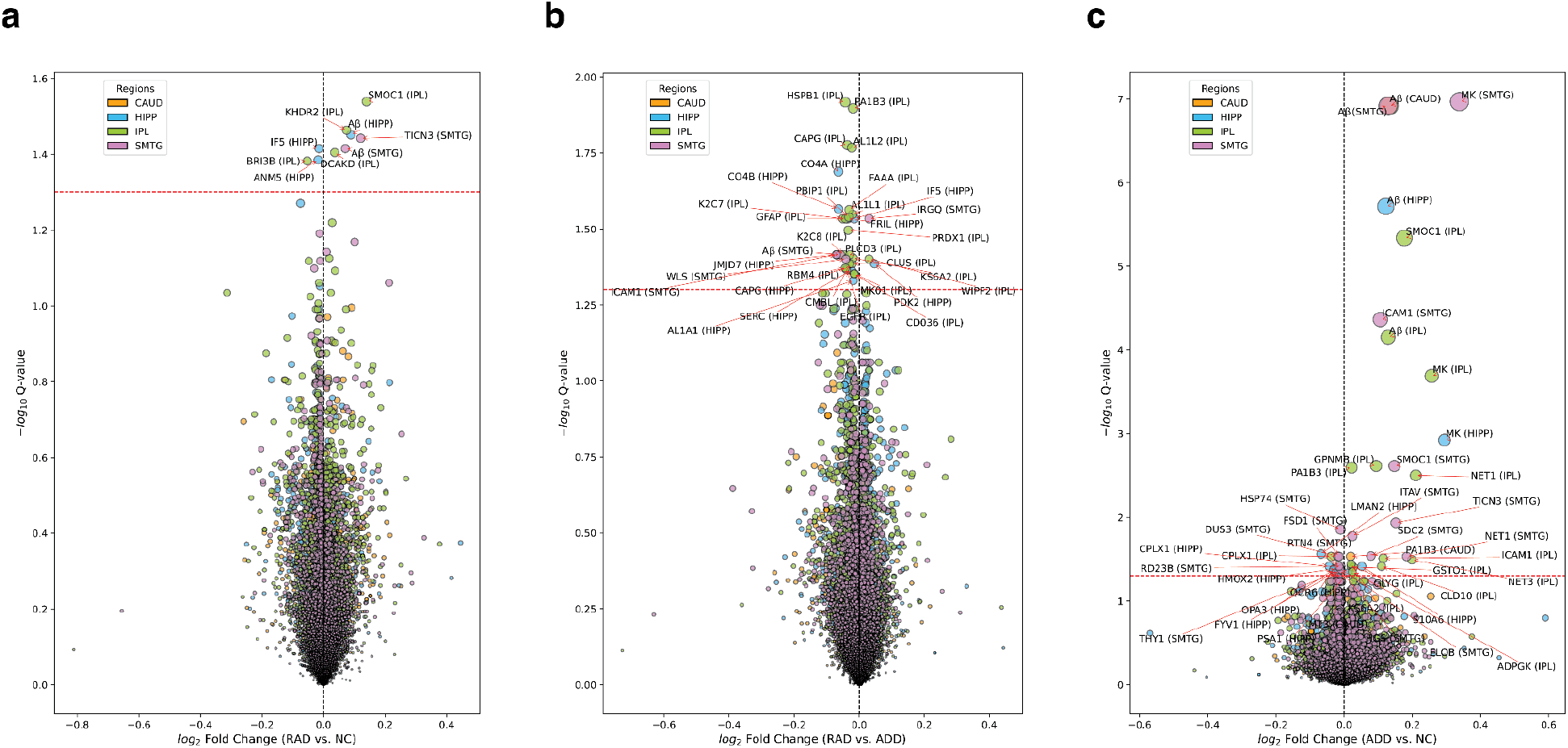
Volcano plot of proteins from four brain regions for (a) RAD vs. NC, (b) RAD vs. ADD, and (c) ADD vs. NC. The dashed red lines are adjusted P-value = 0.05. Circle size is -log_10_(adjusted P-value). Most proteins were differentially expressed between RAD vs. ADD and NC vs. ADD. Most significantly differentially expressed proteins between RAD vs. ADD were lower in the RAD group. Most significantly differentially expressed proteins between NC and ADD were lower in the NC group.

**Extended Figure 3.**
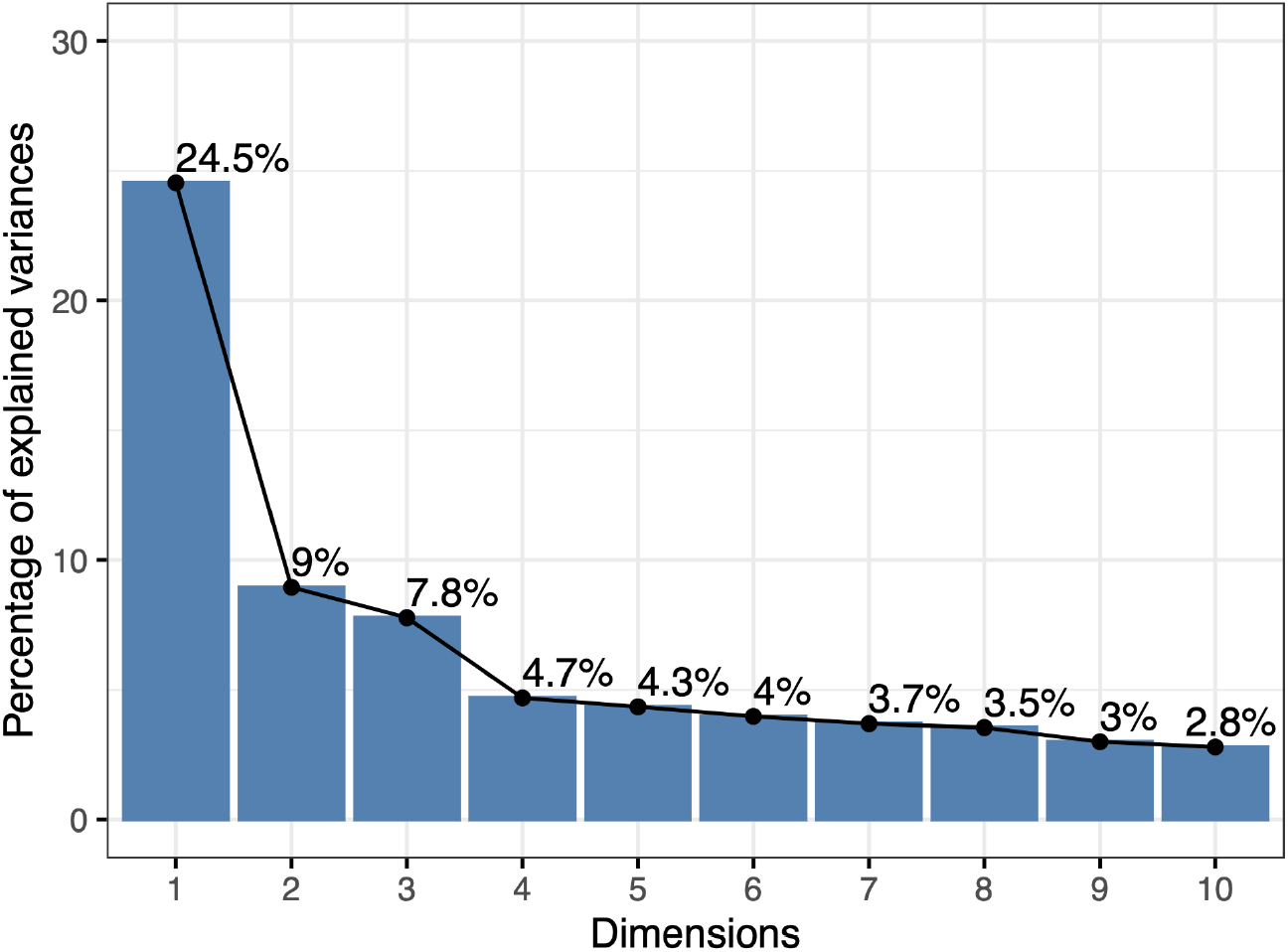
Percentage of explained variances from principal component analysis (PCA) performed with 33 RAD DEPs in all 4 brain regions (original dimension: 33 × 4 = 132).

**Extended Figure 4.**
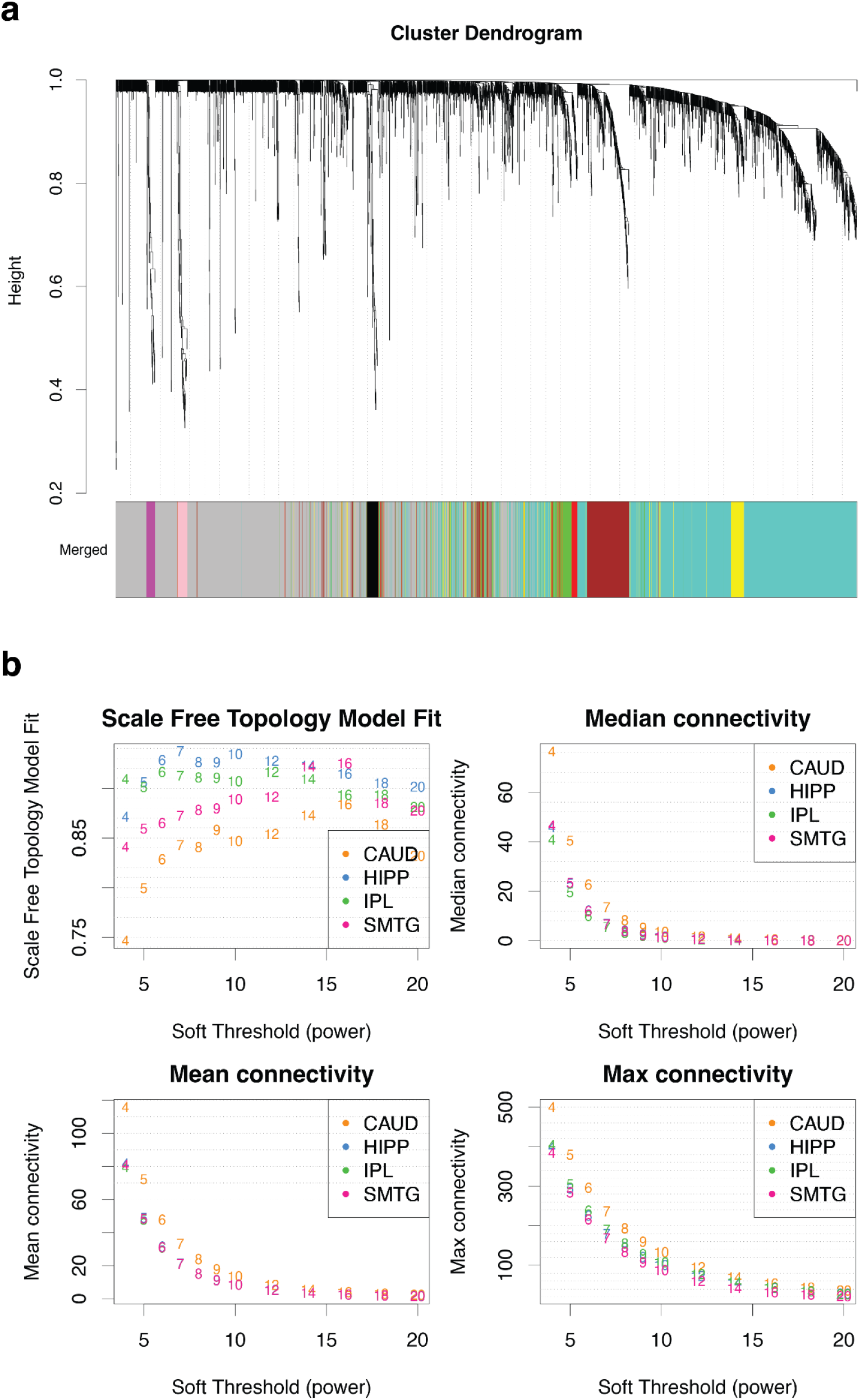
(a) WGCNA co-expression network cluster dendrogram. (b) WGCNA soft threshold vs. scale-free topology, median, mean, and max connectivity.

**Extended Figure 5.**
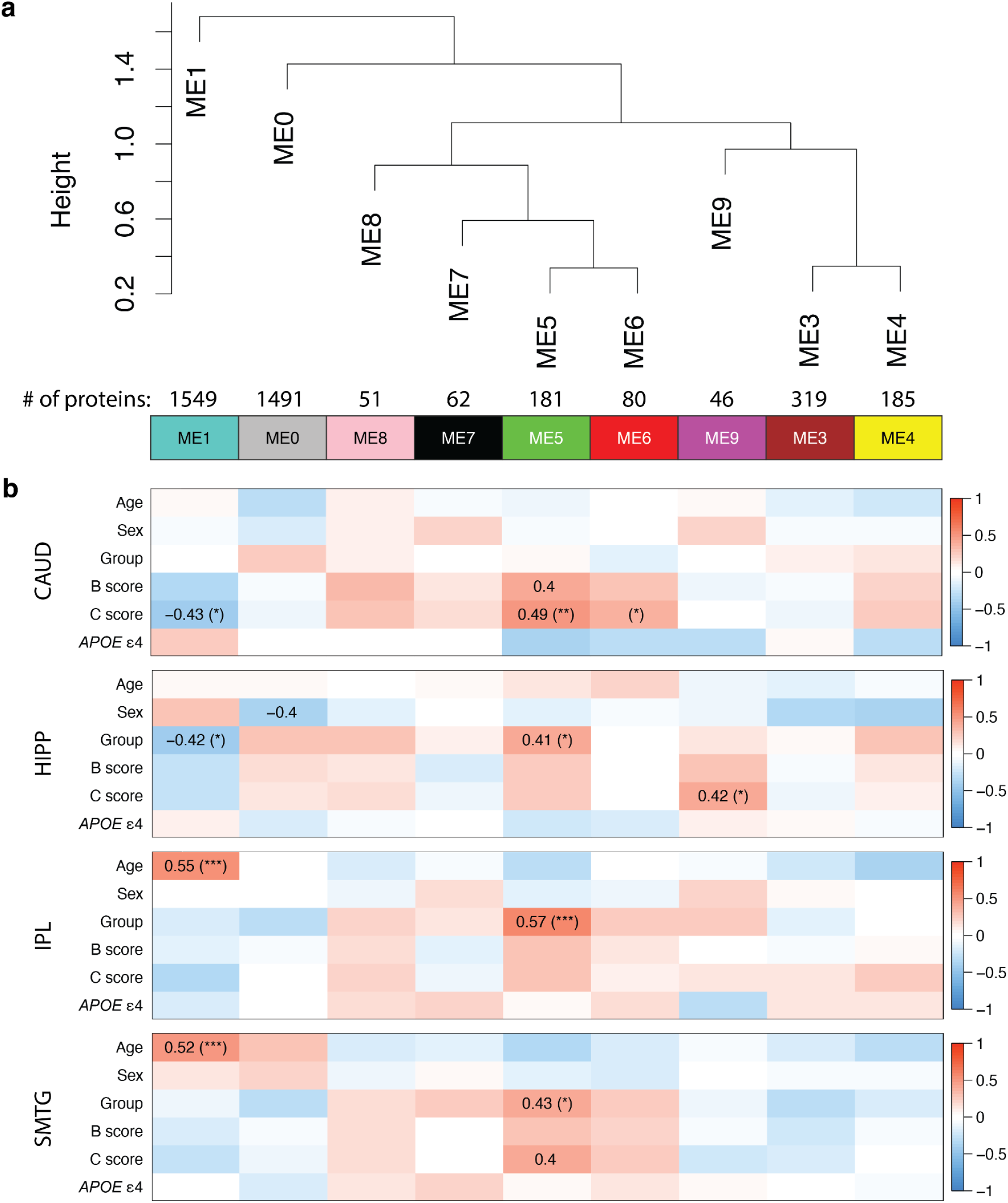
(a) Consensus protein co-expression analysis developed 9 modules across four brain regions. (b) Bi-weighted mid-correlation was used to evaluate the relationships between different features and eigenprotein expression (correlation text threshold: ±0.4). All P-values were adjusted by the Benjamini-Hochberg procedure (FDR cut-off = 0.05).

**Extended Figure 6.**
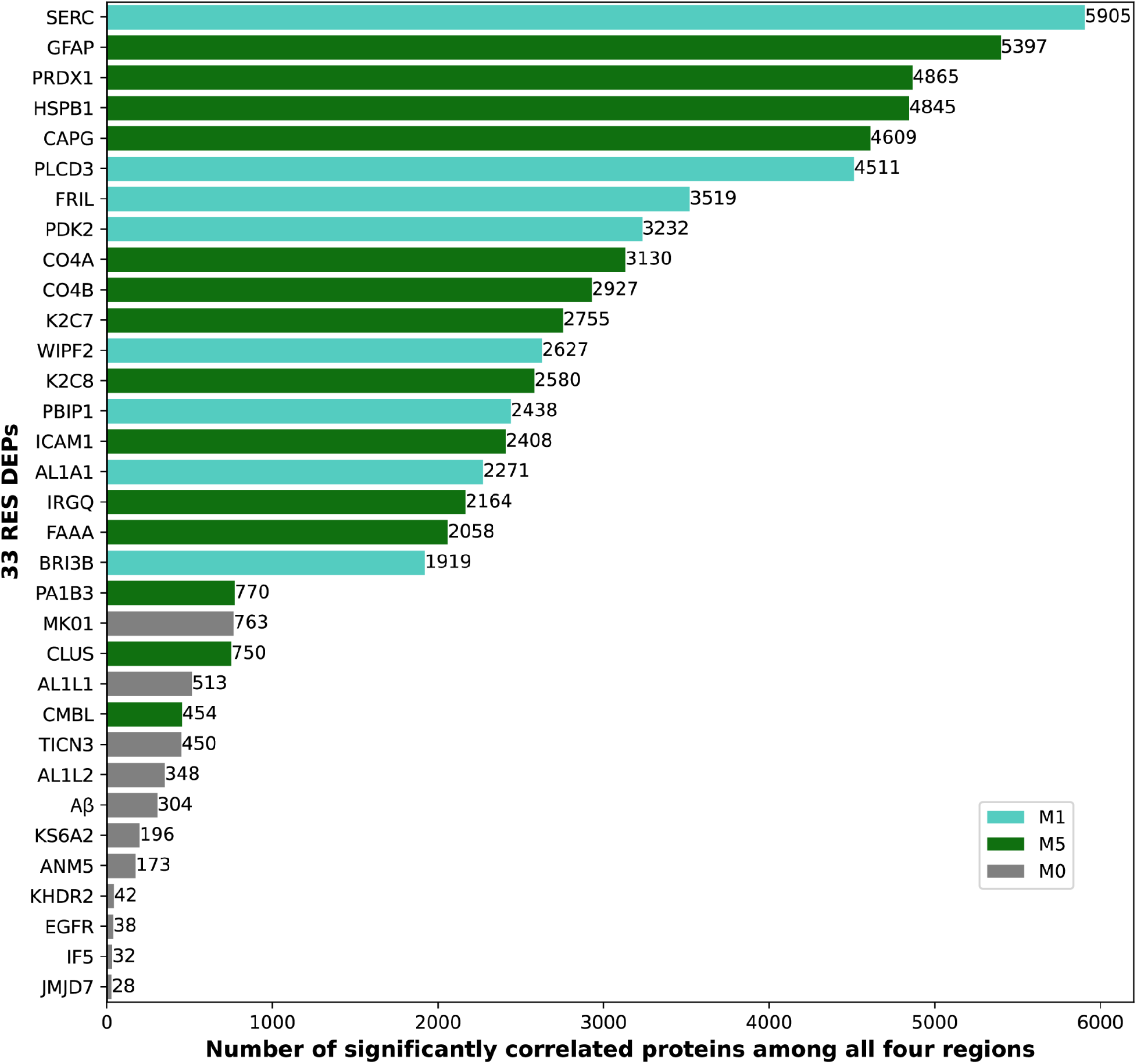
The 33 RAD DEPs were sorted in descending order based on the number of co-expressed proteins (Spearman adjusted P-value < 0.05). Proteins that were not commonly co-expressed with others were more likely to be attributed to M0 (unassigned module).

**Extended Figure 7.**
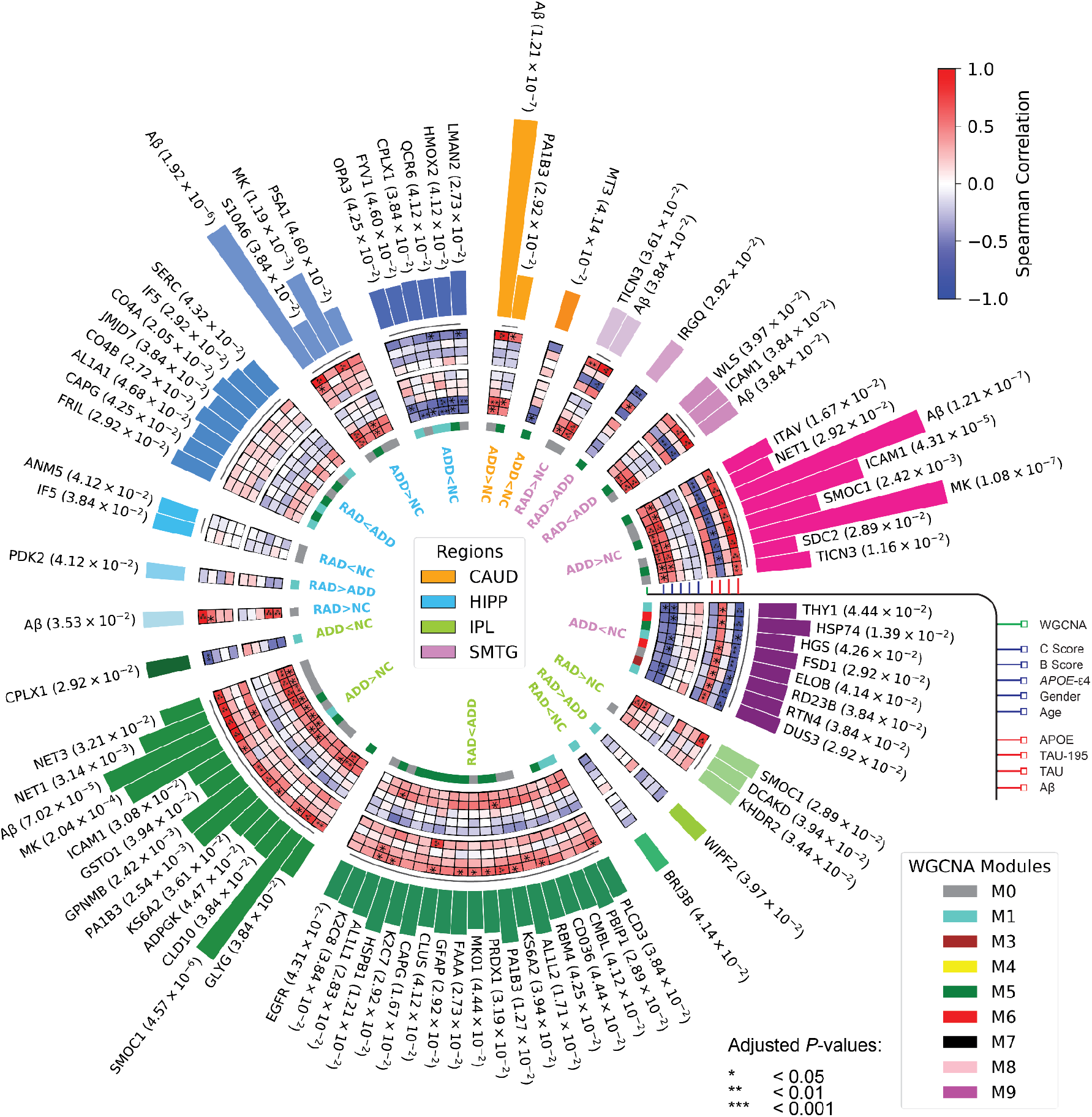
85 differentially expressed proteins in at least one of the four brain regions. Comparisons include RAD vs. NC, RAD vs. ADD, and ADD vs. NC. Circles from outer rings to inner rings: (i) two-sided Student’s t-test adjusted p-value; (ii) correlations with four hallmark proteins; (iii) correlations with individual features; and (iv) associated WGCNA modules.

**Extended Figure 8.**
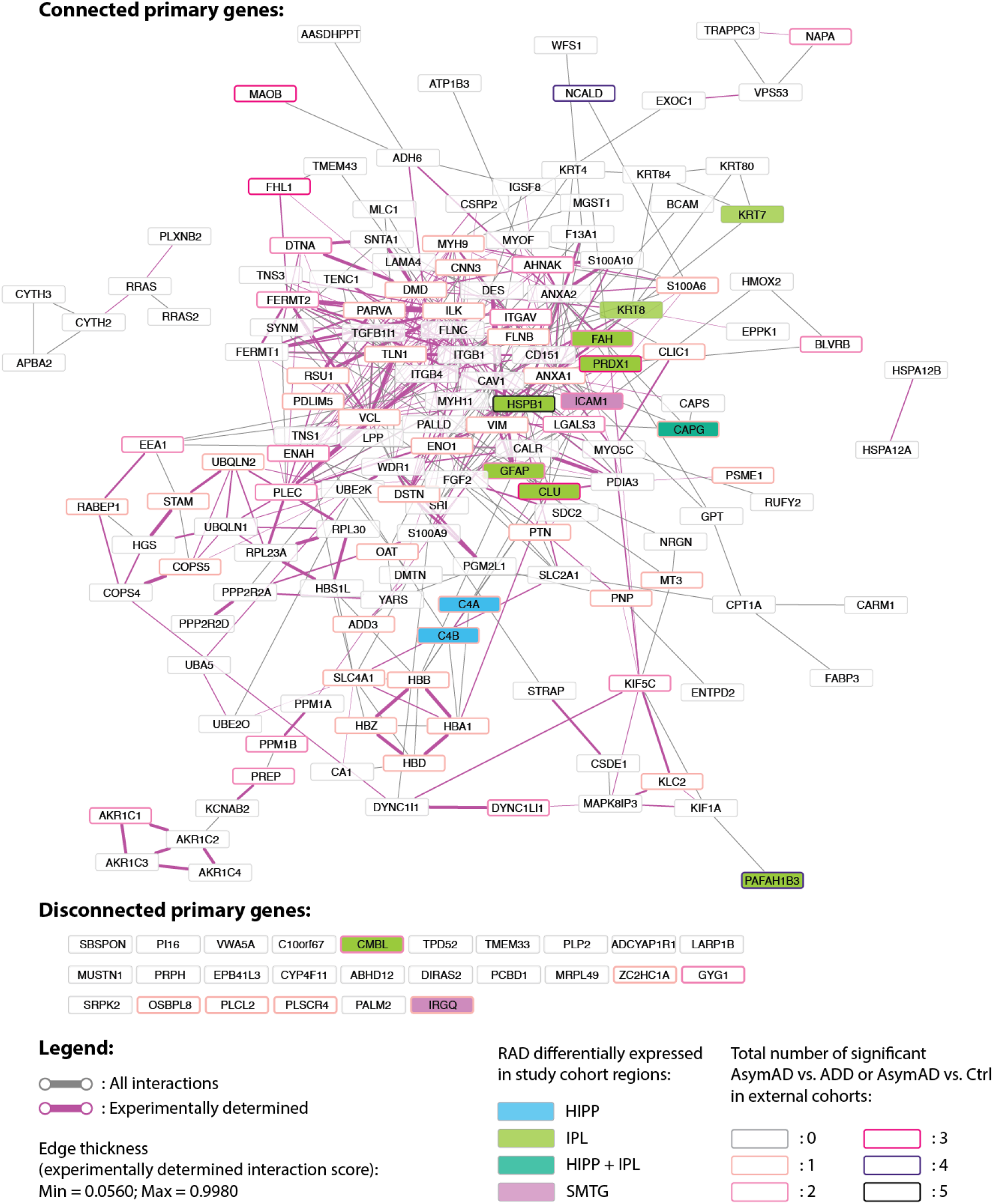
Protein-protein interactions of module 5. Minimum required interaction combined score = 0.4 (default). Magenta-colored edges represent experimentally determined interactions, and thicknesses of edges indicate the experimentally determined interaction score (Min=0.056; Max=0.998). 14 RAD DEPs were highlighted and colored by brain region. Significant differences between AsymAD vs. ADD or AsymAD vs. Ctrl in external datasets were emphasized by thicker borders.

**Extended Figure 9.**
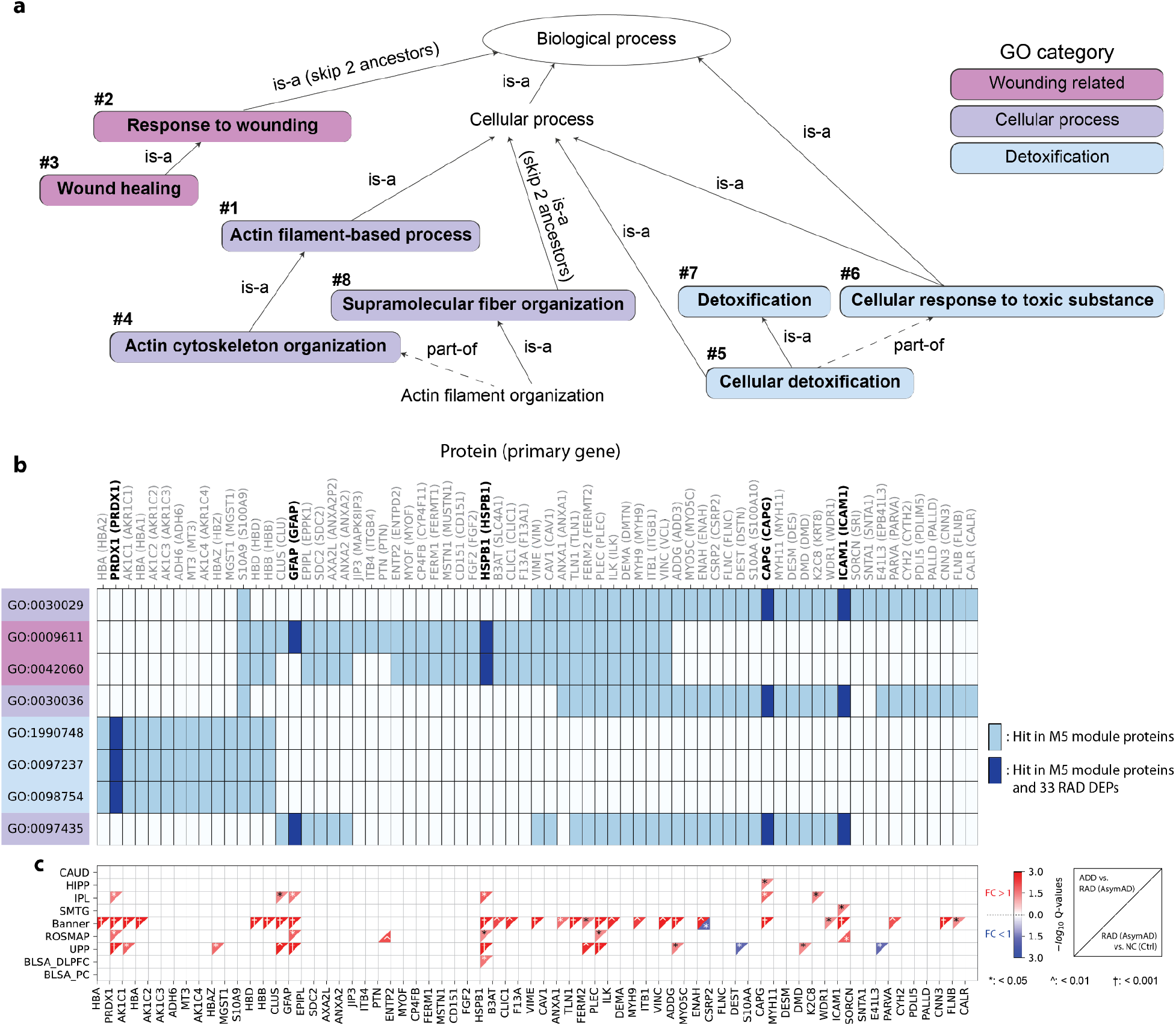
(a) Ancestor chart of top enriched GO biological processes in module 5. (b) Top 8 enriched GO biological processes with proteins hit in the M5 module. (c) Corresponding differential expression analysis results on ADD vs. RAD (or AsymAD) and RAD (or AsymAD) vs. NC (or Ctrl).

**Extended Figure 10.**
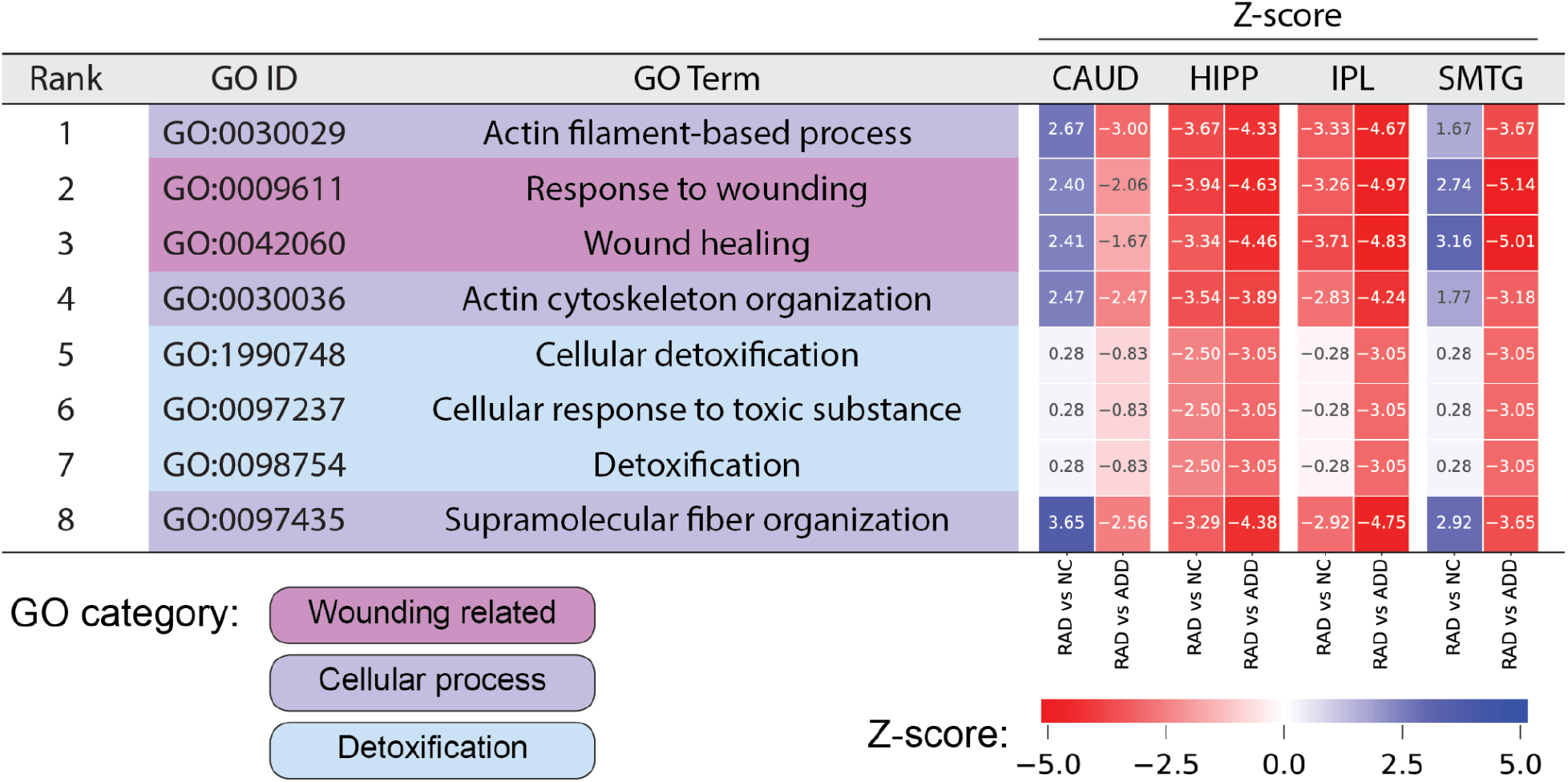
Top 3 enriched GO biological process categories in M5 and their z-scores in study cohorts. Z-scores were calculated based on the number of up/down regulated proteins from hit count in the query, providing an overall pattern between RAD group and GO terms. A negative z-score means that more than half of the proteins had lower expression in RAD than NC or in RAD than ADD (fold-change < 1).

**Extended Figure 11.**
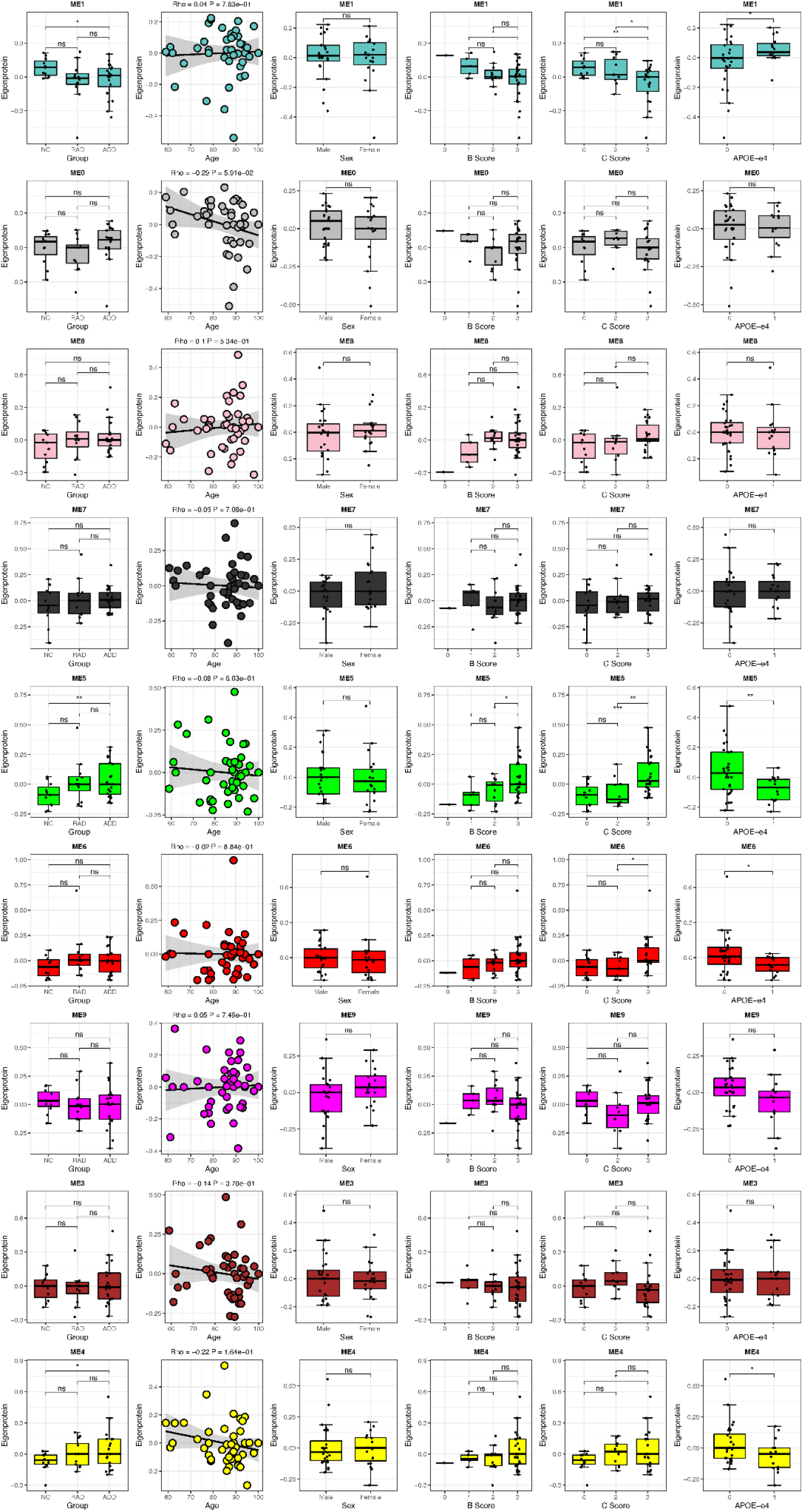
Eigenprotein expressions versus group, age, sex, B score, C score, and *APOE* ε4 allele in CAUD.

**Extended Figure 12.**
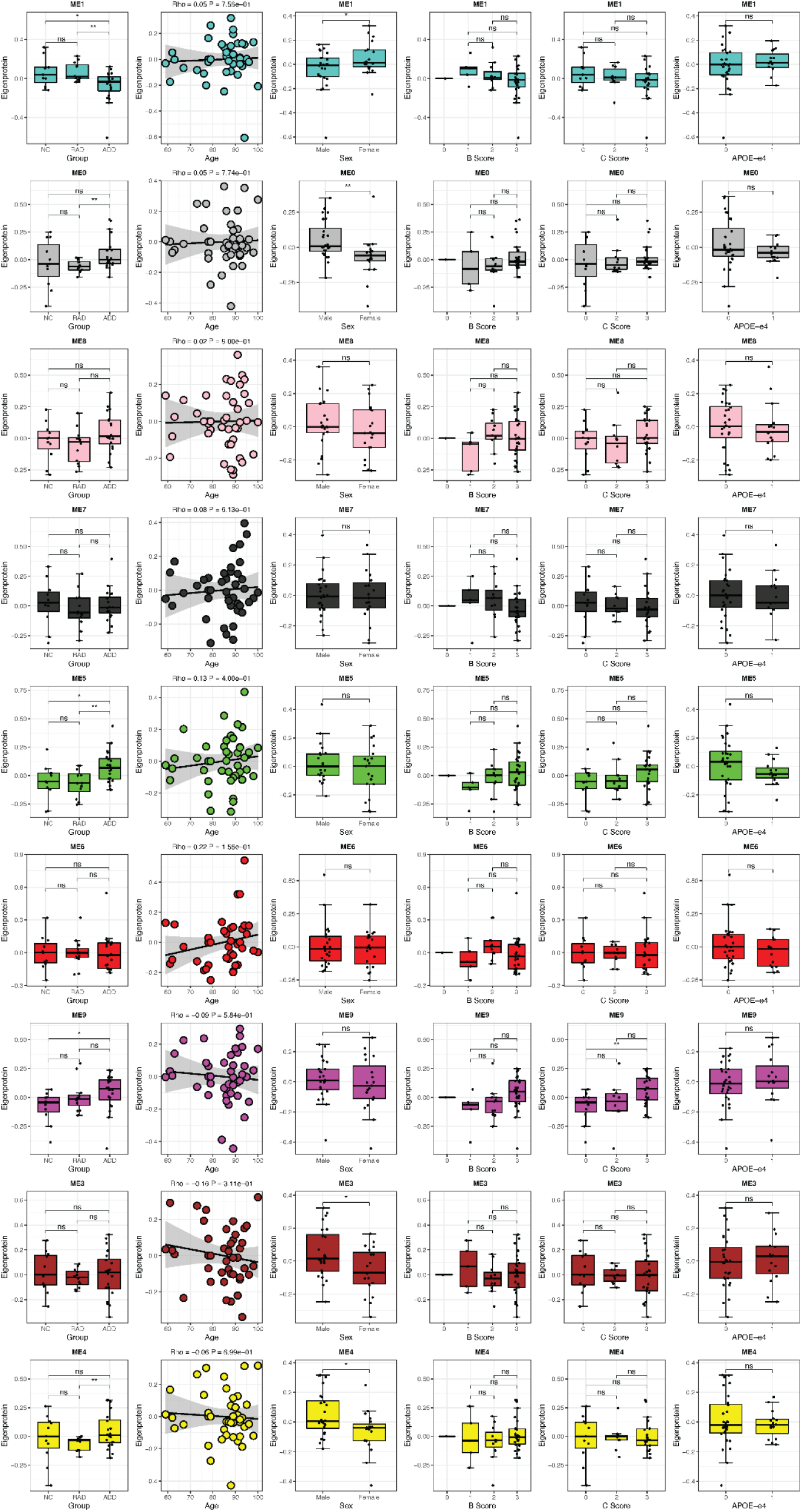
Eigenprotein expressions versus group, age, sex, B score, C score, and *APOE* ε4 allele in HIPP.

**Extended Figure 13.**
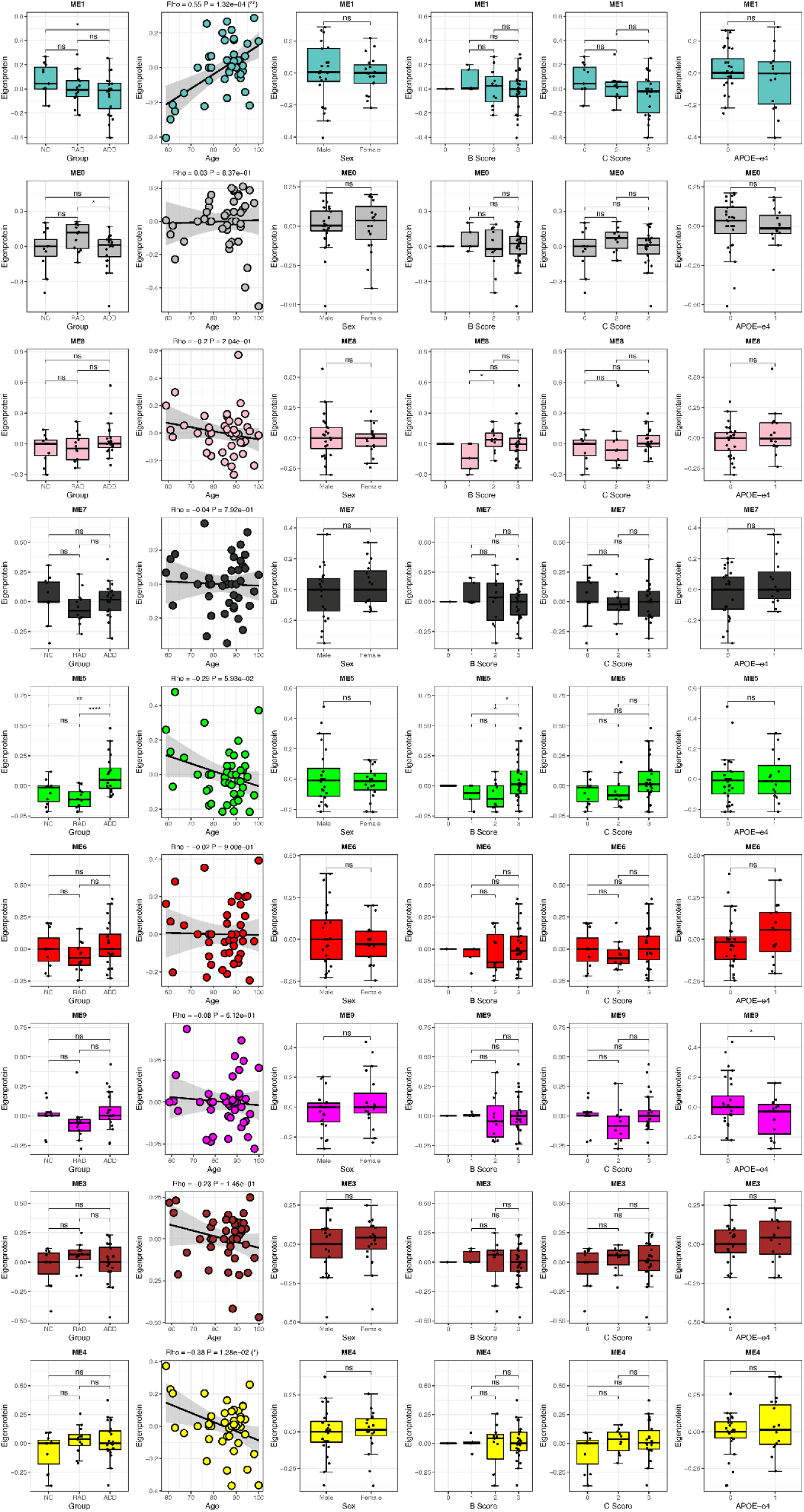
Eigenprotein expressions versus group, age, sex, B score, C score, and *APOE* ε4 allele in IPL.

**Extended Figure 14.**
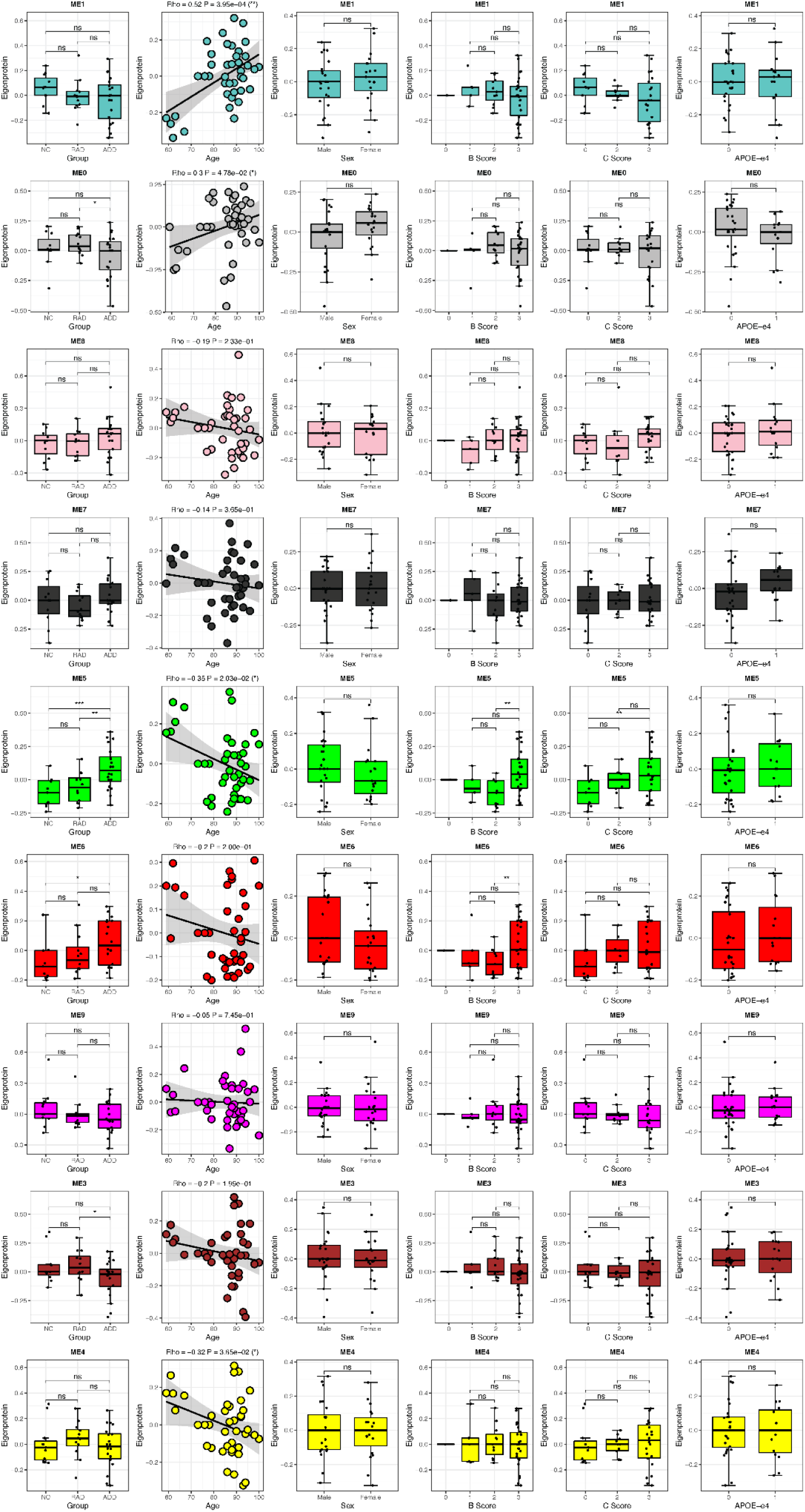
Eigenprotein expressions versus group, age, sex, B score, C score, and *APOE* ε4 allele in SMTG.

**Extended Figure 15.**
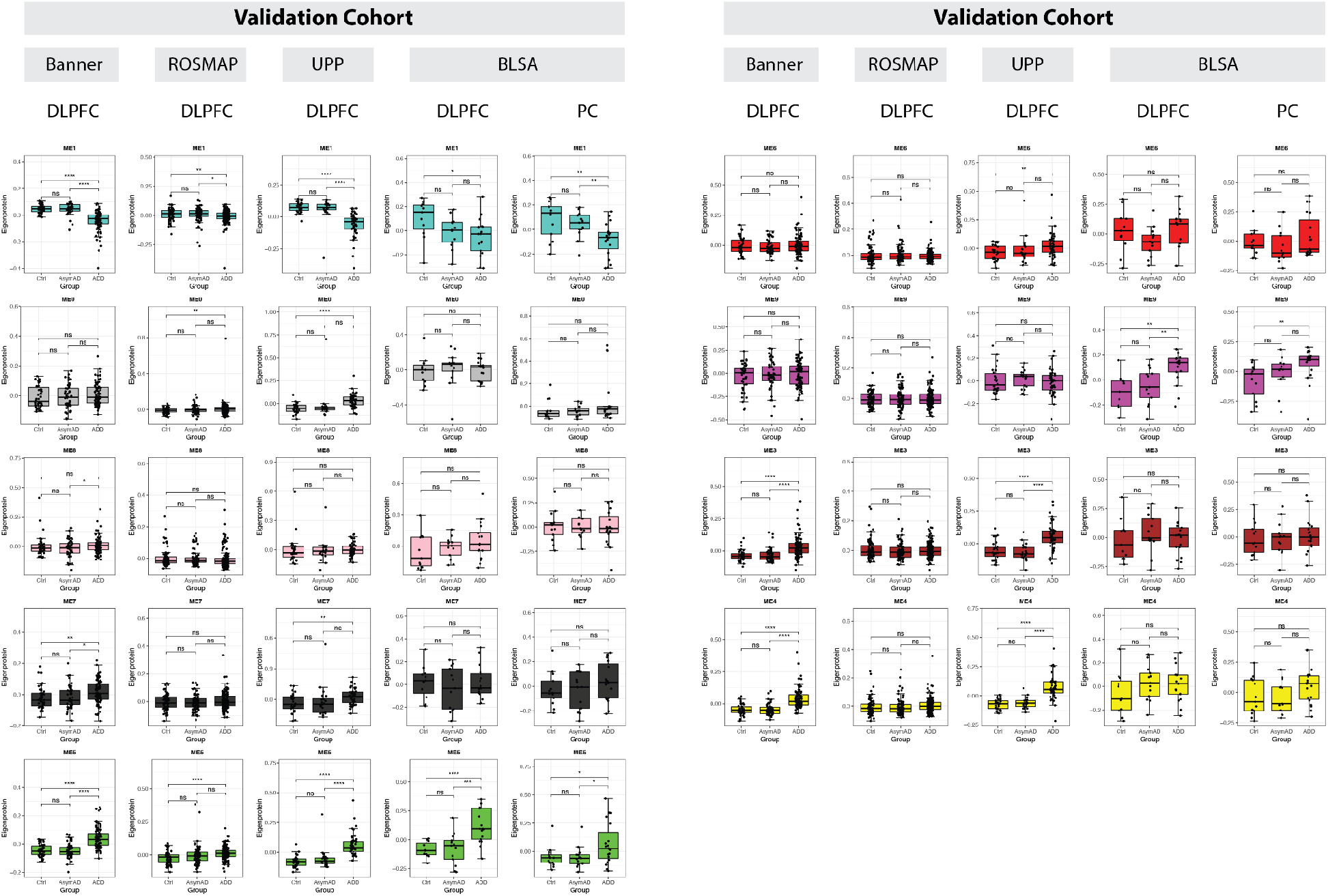
Nine co-expression modules from the study cohort were validated by external datasets. AsymAD is a mix of preclinical AD and RAD cases. Ctrl includes NC.

